# RIG-I-like receptors sense replication stress through ribosomal DNA integrity to drive inflammation

**DOI:** 10.64898/2026.05.27.728122

**Authors:** I-Hui Wu, Xi Wang, Ryan Clark, Soroor Farahnak, Sadeem Ahmad, Sun Hur

## Abstract

Replication stress, a major source of DNA damage and vulnerability in cancer, activates innate immunity through both the cGAS–STING and RIG-I–like receptor (RLR) pathways, yet the mechanism for RLR activation remains poorly understood despite frequent suppression of cGAS–STING in cancer. Here, we show that replication stress induces genomic instability at ribosomal DNA (rDNA), accompanied by aberrant RNA polymerase I–dependent sense and antisense transcription that generates immunostimulatory RNA activating RLR signaling. RLR activation required CDK1 activity but occurred independently of mitosis or micronuclei formation. Cas9-mediated disruption of transcribed rDNA regions was sufficient to induce dysregulated rDNA transcription and RLR activation. Human tumor genomes also showed increased structural variations at rDNA, consistent with rDNA instability. Together, these findings reveal how genomic instability drives inflammation through rDNA-derived immunostimulatory RNA and identify a previously unappreciated consequence of damage-induced transcription.

## Introduction

Self–nonself nucleic acid discrimination is fundamental to all immune systems from bacteria to humans. In vertebrates, the RIG-I-like receptors (RLRs), RIG-I and MDA5, survey cytoplasmic RNA for features characteristic of viral origin, while cGAS detects cytosolic DNA^1–3^. These RNA- and DNA-sensing pathways converge on shared downstream signaling modules, including activation of the transcription factor IRF3 and induction of type I interferons, which in turn elicit a second wave of transcriptional activation of hundreds of antiviral genes via STAT1/2 transcription factors (Fig. 1A). RIG-I detects viral double-stranded RNAs (dsRNAs) bearing 5′-triphosphates (5’ppp) or -diphosphates generated by many negative strand RNA viruses, whereas MDA5 recognizes long dsRNA replication intermediates often accumulated by positive-strand RNA viruses^1^. Increasing evidence indicates that RLRs and cGAS can also be activated by endogenous nucleic acids generated under various conditions of cellular dysregulation, contributing to “sterile” inflammation that is now recognized as a basis for a variety of pathologies, from aging to neurodegenerative diseases^1–3^.

**Fig. 1.**
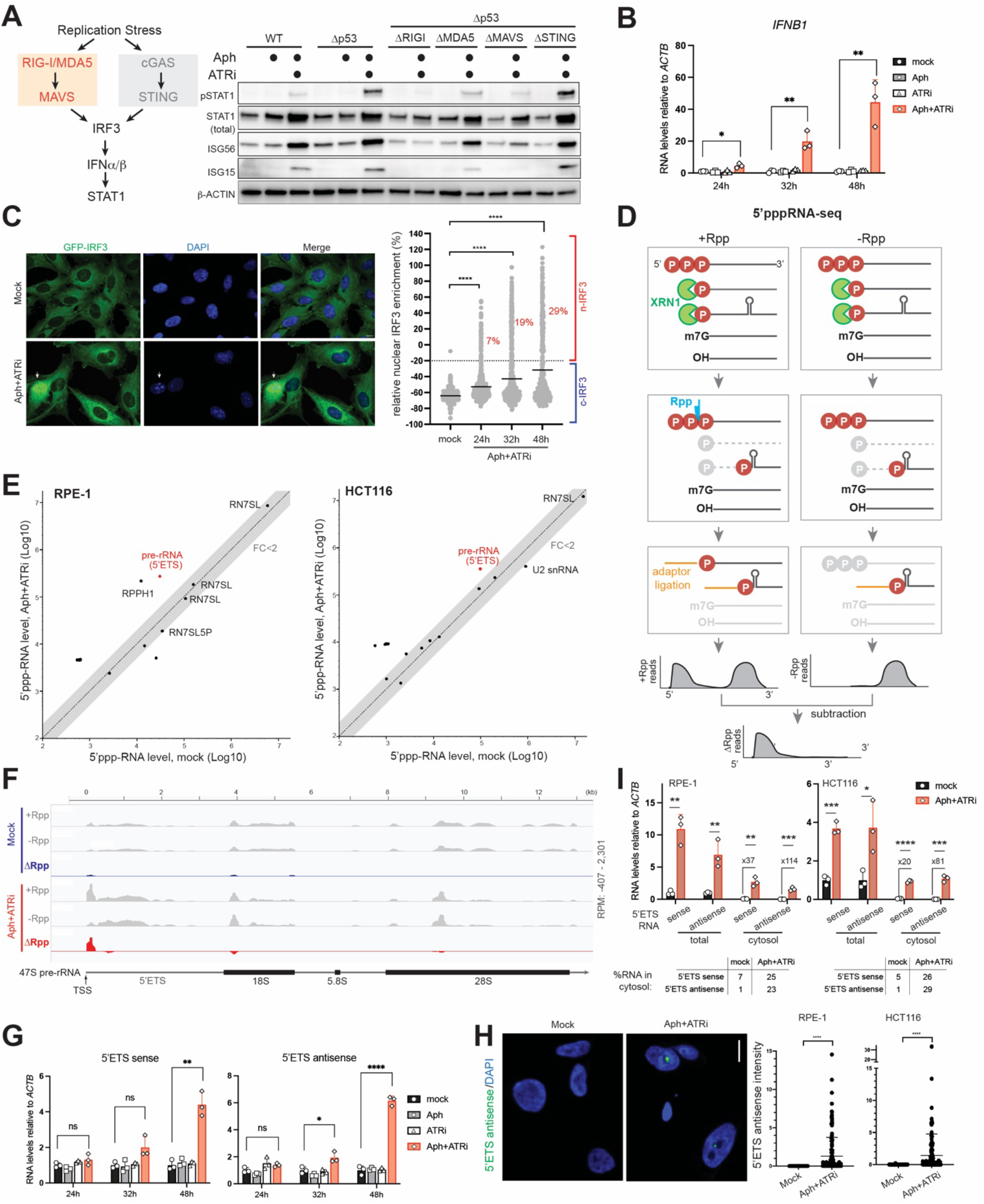
Replication stress induces aberrant rDNA transcription in sense and antisense orientations. (A) Schematic of innate immune signaling pathways activated by replication stress (left) and western blot analysis of STAT1 phosphorylation (pSTAT1), induction of interferon-stimulated genes (ISG56 and ISG15) (right). RPE-1 cells of indicated genotypes were treated with aphidicolin (Aph, 0.5 μM) and ATR inhibitor (VE-821; ATRi, 2.5 μM) for 48 h prior to analysis. β-ACTIN was used as a loading control. (B) RT-qPCR analysis showing levels of *IFNB* mRNA relative to *ACTB* 24, 32 and 48h post Aph+ATRi-treatment in RPE-1 cells. Values represent mean ± SD from 3 biological repeats. \*\**p* <0.01, \**p* <0.05 from two-tailed student’s t test. (C) IRF3 nuclear translocation upon Aph+ATRi treatment. Left: Representative immunofluorescence images of lentivirally expressing GFP-IRF3 (green) and nuclei (DAPI, blue) in RPE-1 cells under mock condition or Aph+ATRi treatment for 24 h. White arrows indicate cells with nuclear IRF3. Scale bar, 10 μm. Right: Quantification of nuclear IRF3 enrichment (%), as defined by the difference between mean nuclear and mean cytoplasmic GFP-IRF3 intensity, normalized to the mean cytoplasmic intensity. Each data point represents an individual cell (n=399, 596, 504, 341 from left to right) from ~60 fields of view per condition. Cells with >-20% nuclear enrichment were classified as nuclear IRF3 (n-IRF3) cells. The rest were classified as cytosolic IRF3 (c-IRF3) cells. Percentages of n-IRF3 cells at indicated time points are shown in red. \*\*\*\**p* < 0.0001 by two-tailed Mann–Whitney test. (D) 5′pppRNA-seq workflow: Pre-existing 5′p-RNAs are first degraded by XRN1, followed by RNA 5’ polyphosphatase (Rpp) treatment to convert 5′ppp/5′pp ends into 5′p termini for adaptor ligation. 5′ppp-containing RNAs are then identified as Rpp-enriched regions by subtracting −Rpp from +Rpp coverage (ΔRpp), followed by peak calling on the ΔRpp track. (E) Comparison of 5’ppp-RNA levels between mock and Aph+ATRi-treated cells (RPE-1 and HCT116). Each point represents an 5’ppp-RNA species identified by ΔRpp peak calling and is plotted using normalized AUC (area under curve) values in mock vs. Aph+ATRi-treated cells. The dashed line denotes equal abundance between conditions, and the shaded region indicates fold change (FC) upon Aph+ATRi treatment <2. Two biological repeats were used in all analyses. Multiple data points for RNA7SL reflect mapping to the Repbase reference and to multiple genomic loci in the hg38 reference through tiered mapping (See Methods). (F) Representative IGV tracks of 5′ppp-RNA-seq across the 47S pre-rRNA locus in mock and Aph+ATRi-treated RPE-1 cells. The +Rpp and −Rpp tracks display multiple prominent peaks across the pre-rRNA, likely due to incomplete RNA digestion by XRN1. In contrast, the ΔRpp track with Aph+ATRi only shows one prominent peak corresponding to 5’ETS and beginning at the known transcription start site (TSS) of pre-rRNA. Bottom: 47S pre-rRNA schematic indicating 5′ETS, 18S, 5.8S, and 28S. (G) RT-qPCR analysis of 5’ETS sense and antisense RNAs relative to *ACTB* 24, 32 and 48h post Aph+ATRi-treatment in RPE-1 cells. Expression levels are normalized against the corresponding mock levels. Values represent mean ± SD from 3 biological repeats. \*\*\*\**p* <0.0001, \*\**p* <0.01, \**p* <0.05, not significant (ns) *p* > 0.05 from two-tailed student’s t test. (H) RNA-FISH analysis showing antisense 5’ETS in RPE-1 and HCT116 cells upon mock and Aph+ATRi treatment for 48 h. Antisense probes map to the rDNA locus −1080 to +241 relative to TSS. Each data point represents an individual cell (n=382 and 142 for mock and Aph+ATRi RPE-1; n=54 and 146 for mock and Aph+ATRi HCT116) from ~20 z-stacked fields of views. ****, *p*<0.0001 by a two-tailed nonparametric Mann–Whitney–Wilcoxon test. Images on the left are RPE-1 cells. Scale bar, 10 µm. (I) RT-qPCR analysis showing levels of 5’ETS sense and antisense RNAs relative to *ACTB* 48h post Aph+ATRi-treatment in RPE-1 and HCT116 cells. Expression levels are plotted relative to mock (total) for sense and antisense separately. The table below shows the percentage of 5’ETS sense and antisense RNAs localized to cytosol under mock and Aph+ATRi condition. Values represent mean ± SD from 3 biological repeats. \*\*\*\**p* <0.0001, \*\*\**p* <0.001, \*\**p* <0.01, \**p* <0.05, from two-tailed student’s t test.

One such condition is replication stress, which arises when DNA replication is impeded by obstacles such as transcription– replication conflicts, depletion of nucleotide pools, or unscheduled origin firing, and often leads to DNA damage^4,5^. Replication stress is a pervasive feature of cancer due to the high proliferative demand of tumor cells, and is therapeutically exploited by chemo- and radiotherapies, which further exacerbate replication stress to suppress tumor growth and to promote anti-tumor innate immune responses^6–8^. While most prior work has focused on cGAS activation in these settings^2,9–12^, studies have suggested that RLRs also play an important, and in some cases dominant role^13–16^, particularly in tumors where cGAS–STING signaling is suppressed^17–19^. However, how RLRs are activated under replication stress remains unclear, including the identity and genomic origin of the endogenous RNA ligands involved.

Ribosomal DNA (rDNA) repeats are a unique class of genomic elements. In humans, rDNA is present in several hundred copies, clustered across five distinct chromosomes (chr 13, 14, 15, 21, 22)^20,21^. Each repeat unit (~43 kb) harbors an intergenic, untranscribed spacer (IGS, ~30 kb) and a region transcribed by RNA polymerase I (RNAP I) as a single precursor pre-rRNA (47S, 13.3 kb), which is subsequently processed into mature rRNAs (18S, 5.8S and 28S). Among the hundreds of copies of rDNA, which vary across individuals, only a fraction of them are thought to be actively transcribed at any given time, and these repeats cluster within the nucleolus, where pre-rRNA is synthesized and processed into ribosomes^20,21^. Despite the ribosome being one of the best-characterized cellular machines, rDNA itself remains among the least well-characterized regions of the human genome and is commonly excluded from most functional genomic analyses.

Here, we show that replication stress induces rDNA instability, and this results in aberrant bidirectional transcription of rDNA and accumulation of 5’ppp-containing dsRNA that drives RLR activation. We further show that rDNA break alone is sufficient to activate RLRs and is commonly observed in human tumors. Together, these findings define the mechanism by which replication stress links DNA damage to RNA-based innate immunity.

### Replication stress induces aberrant rDNA transcription in sense and antisense orientations

To investigate how replication stress triggers RLR activation, we used hTERT-immortalized non-cancerous human retinal pigment epithelial cells (RPE-1) and colorectal carcinoma cells (HCT116), both of which are known to lack the functional cGAS-STING pathway^13,14^. Replication stress was induced by low-dose aphidicolin (Aph, 0.5 μM), a well-established condition for replication stress^13,22^. Previous studies^13,14^ have shown that ATR inhibition, an emerging therapeutic strategy for a wide range of cancers, together with p53 deficiency, which is common across many tumor types, accelerates and amplifies the innate immune response. We therefore used this sensitized setting to enable robust detection and mechanistic analysis of early signaling events. Consistent with prior reports^13,14^, antiviral signaling, as measured by phosphorylated STAT1 (pSTAT1) and interferon-stimulated genes (e.g., STAT1 (total), ISG15, ISG56), was significantly enhanced at day 2 when Aph was combined with ATR inhibitor (VE-821; ATRi; 2.5 μM,) in p53-deficient backgrounds (Δp53, Fig. 1A and fig. S1A). Aph+ATRi–induced signaling depended on RIG-I, MDA5 and their downstream adaptor MAVS, but not STING, in both RPE-1 and HCT116 cells (Fig. 1A and fig. S1, A and B), consistent with previous reports that cGAS-STING signaling is inactive in these cells^13,14^.

Aph+ATRi–induced innate immune signaling was also evident from *IFNB1* transcriptional induction (Fig. 1B) and nuclear translocation of the upstream transcription factor IRF3 (Fig. 1C). All three readouts—IRF3 translocation (Fig. 1C), *IFNB1* mRNA (Fig. 1B) and STAT1 phosphorylation (fig. S1C)—showed a gradual increase in immune signaling over 32-48 h of Aph+ATRi treatment. Because IRF3 nuclear translocation represents an upstream event that directly reflects RLR activation, independent of paracrine IFN signaling, we next quantified the fraction of cells activated. ~20–30% of cells exhibited nuclear IRF3 (n-IRF3) at 32-48 h (Fig. 1C). Overexpression of RIG-I did not significantly increase the fraction of n-IRF3 cells (fig. S1D), suggesting that incomplete penetrance of IRF3 activation is not due to limiting RIG-I levels, but instead reflects other constraints, possibly the availability of endogenous RNA ligands for RLRs.

We next examined how RLRs are activated upon replication stress and which endogenous RNA ligands are induced by Aph+ATRi. Given that RIG-I deletion had more significant impact on pSTAT1 level than MDA5 deletion (Fig. 1A, fig. S1A), we started with RIG-I investigation. RIG-I typically recognizes 5′-triphosphorylated or diphosphorylated (5′ppp/5′pp) dsRNA, but can also respond to RNAs with noncanonical caps (e.g., FAD-, NAD-, or incompletely methylated 7mG caps)^23^. To examine whether endogenous RIG-I ligands under replication stress harbor 5’ppp/5’pp or non-canonical caps, we utilized DUSP11, a phosphatase that removes 5′ppp/5′pp but not noncanonical caps^24^. DUSP11 knockout amplified Aph+ATRi-induced antiviral signaling, whereas catalytic DUSP11 overexpression suppressed it (fig. S1E). Overexpression of catalytically dead variant of DUSP11 (mDUSP11) did not suppress antiviral signaling (fig. S1E). These results suggest that RIG-I is activated by endogenous RNA bearing 5′ppp/5′pp ends during replication stress.

To identify the RNA species responsible for RIG-I activation, we adapted and optimized a previously published 5′pppRNA-seq method^25,26^, using the bacterial polyphosphatase Rpp to convert 5′ppp/5’pp into 5′p suitable for sequencing adaptor ligation, after removal of pre-existing 5′p-RNA with 5’p-end-specific exoribonuclease XRN1 (Fig. 1D). XRN1 digestion can be incomplete, especially in structured RNA regions, which would leave behind a 5’p-end on the degradation intermediate that would contribute to a background signal (Fig. 1D). To distinguish true 5’ppp-containing RNAs from this background, we leveraged the fact that adaptor ligation of 5’ppp-RNA requires prior Rpp treatment, whereas XRN1-derived background does not. We therefore performed sequencing with and without the Rpp treatment and calculated differential coverage (ΔRpp) to identify Rpp-dependent 5’ppp-RNA species (Fig. 1D).

Consistent with the notion that Rpp-sensitive sites contain a 5’ppp-end, RNA known to harbor 5’ppp, such as the RNA polymerase III (RNAP III) transcripts 7SL RNA and H1 RNA subunit of RNase P (RPPH1), showed strong ΔRpp enrichment (fig. S2A). In contrast, RNAP II transcripts, such as *GAPDH* and *ACTB* mRNAs, which carry a 7-methylguanosine cap at their 5’ ends showed little to no ΔRpp enrichment (fig. S2B). Certain internal regions showed coverage in the +Rpp condition (fig. S2B), but comparable signal was observed in the −Rpp control, resulting in no peak in ΔRpp coverage.

Systematic comparison of ΔRpp signals between mock vs. Aph+ATRi-treated cells revealed 47S pre-rRNA as the most prominent 5’ppp-RNA species upregulated by Aph+ATRi-treatment in both RPE-1 and HCT116 cells (Fig. 1E). The 5’ edge of ΔRpp peak mapped precisely to the known RNAP I transcription start site (TSS) and resided within the 5’-external transcribed spacer (5′ETS) upstream of the mature rRNAs regions (Fig. 1F, and fig. S2C). Under basal condition, 5’ETS would be rapidly removed from pre-rRNA and cleared through a combination of endonucleolytic cleavage and exonucleolytic digestion^27^. RT-qPCR across cleavage junctions indicated that pre-rRNA processing was compromised in the presence of Aph+ATRi (fig. S2D). Nanopore sequencing revealed that

5′ppp-containing pre-rRNAs were predominantly ~230 nt in length (fig. S2E), well upstream of the first cleavage site A’/01 (at position ~415), suggesting abortive rDNA transcription and/or processing in Aph+ATRi-treated cells.

In support of aberrant transcriptional activity, strand-specific RT-qPCR showed that antisense transcription also occurred at 5’ETS in response to Aph+ATRi (Fig. 1G). RNA-FISH analysis indicated that antisense 5’ETS RNA was produced within or adjacent to nucleoli, where rDNA is concentrated (Fig. 1H). Notably, the induction was heterogeneous across cells, with a subset showing orders-of-magnitude increases, whereas the majority exhibited little to no change, similar to heterogeneity observed in IRF3 nuclear translocation (Fig. 1C). Both sense and antisense 5’ETS induction occurred over ~32-48 h upon Aph+ATRi treatment and required the combination of Aph and ATRi (Fig. 1G), consistent with kinetics and requirement for *IFNB1* induction (Fig. 1, B and C). Although 5’ETS RNA-FISH signals were more visibly apparent within nucleoli, subcellular fractionation suggests a substantial portion (~23 and 29% in RPE-1 and HCT116 respectively) of these RNAs also localized in the cytoplasm, with ~40-100-fold increase upon Aph+ATRi treatment (Fig. 1I).

Together, these findings indicate that replication stress induces aberrant transcription of both sense and antisense pre-rRNA fragments, including 5’ETS species retaining a 5′ ppp, and that this correlates with RLR signaling.

### Aberrant rDNA transcripts activate RLRs

We next asked whether 5’ETS RNA activates RLRs, and whether this activation requires both sense and antisense RNA. Given the high GC content of 5’ETS (77% in the first 230 nt), it was conceivable that secondary structures within 5’ETS, together with 5’ppp, might be sufficient to activate RIG-I. To test this idea, we performed a well-established *in vitro* signaling assay using purified RNA, recombinant RIG-I, K63-linked ubiquitin (a required cofactor), radiolabeled IRF3, and cell extracts providing downstream signaling components such as MAVS^28,29^ (Fig. 2A, left). In this system, RIG-I activation is measured by IRF3 dimerization on native gels. Upon stimulation with *in vitro-*transcribed dsRNA, robust IRF3 dimerization was observed (Fig. 2A, center). Only the sense-antisense hybrids of 5’ETS, and not either strand alone, activated RIG-I (Fig. 2A, center). Similarly, the hybrid activated MDA5 more efficiently than either strand alone (Fig. 2A, right).

**Fig. 2.**
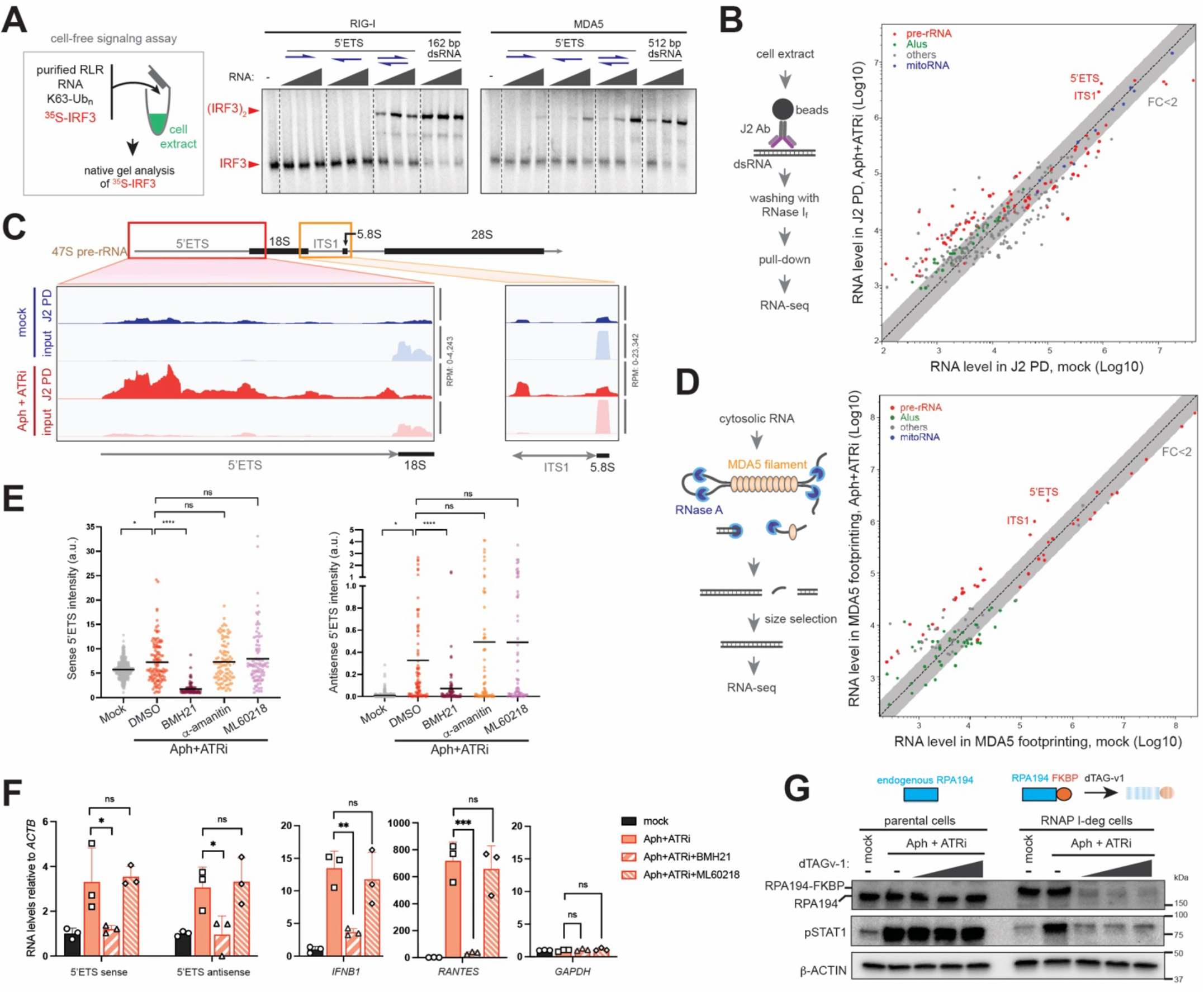
Aberrant rDNA-derived RNAs activate RLRs. (A) Cell-free RLR signaling assay. Left: RLR activation was assessed by monitoring ^35^S-radiolabeled IRF3 dimerization on native gels following incubation with the indicated signaling components and RNA of interest. Center/Right: In vitro transcribed 5’ETS sense and/or antisense RNAs (0.1, 0.5, 2.5 ng/μl) were used to stimulate RIG-I (center) or MDA5 (right). In vitro-transcribed, 5’ppp-containing 162 bp and 512 bp dsRNAs were used as positive control for RIG-I and MDA5 activation respectively. (B) J2 dsRNA pull-down (PD)-seq. Left: Cell extracts from RPE-1 cells treated with Aph+ATRi (48 h) or mock were subjected to J2 PD in the presence of ssRNA-specific nuclease RNase I_f_. Right: Systematic comparison of J2 PD in mock vs. Aph+ATRi-treated cells. Each point represents an RNA species enriched by J2 PD in Aph+ATRi condition, and is plotted using the normalized AUC values from J2 PD-seq in Aph+ATRi vs. mock samples. Data points are colored by RNA class (pre-rRNA, red; Alu, green; mitoRNA, blue; others, gray). Shaded area denotes FC <2 from the comparison of Aph+ATRi vs. mock. (C) Representative IGV tracks showing the 5’ETS and ITS1 regions of pre-rRNA in input vs. J2 PD in mock (blue) and Aph+ATRi-treated samples (red). (D) MDA5 filament footprint-seq assay. Left: Cytosolic RNA from RPE-1 cells were incubated with purified MDA5 protein, followed by RNase A digestion to remove unprotected RNA. RNA was then purified and size-selected to isolate long stretches of dsRNA protected by MDA5 filaments. Right: Systematic comparison of MDA5 footprint-seq in mock vs. Aph+ATRi-treated cells. Each point represents an RNA species enriched by MDA5 footprinting in Aph+ATRi condition, and is plotted using its normalized AUC values from mock vs. Aph+ATRi conditions. (E) RNA-FISH analysis of sense and antisense 5′ETS RNA in RPE-1 cells under mock or Aph+ATRi treatment (48 h). Inhibitors of RNAP I (BMH21, 0.5 μM), RNAP II (α-amanitin, 5 μM), and RNAP III (ML60218, 25 μM) were added 2 h prior to analysis. Each point represents an individual cell (n = 444, 122, 112, 93, 88, 365, 90, 90, 70, 68 from left to right). ~20 fields of view per condition were analyzed. *\*p<0*.*05, ****p<0*.*0001* by two-tailed Mann–Whitney test. (F) RT-qPCR analysis showing RNA levels relative to *ACTB* in RPE-1 cells with or without BMH21 (0.5 μM) or ML60218 (25 μM). Values represent mean ± SD from 3 biological repeats. \*\*\**p*<0.001, \*\**p*<0.01, \**p*<0.05, not significant (ns) *p*>0.05 from one-way ANOVA. (G) Western blot analysis showing RPA194 and pSTAT1 levels in RPE-1Δp53 cells expressing endogenous RPA194 (parental) or degradable RPA194-FKBP in the ΔRPA194 background (RNAP I-deg). Cells were treated with Aph+ATRi with or without dTAG-v1 (0, 0.125, 0.25 and 0.5 μM) for 48 h.

To investigate whether 5’ETS sense and antisense RNAs form duplexes in cells and contribute to RIG-I activation, one approach would be to immunoprecipitate RIG-I and measure the enrichment of 5’ETS RNA. However, we found that this approach does not reliably enrich even the known RIG-I ligands that were exogenously introduced, likely due to RIG-I’s non-specific affinity to non-stimulatory, abundant RNA and its multimerization on stimulatory RNA^30,31^ that would reduce apparent enrichment. Therefore, we instead performed pull-down using dsRNA-specific antibody J2^32^ as an alternative to examine 5’ETS duplex RNA formation. Cell extracts from RPE-1 cells were subjected to immunoprecipitation using J2 in the presence of ssRNA-specific RNase I_f_, followed by RNA-seq (Fig. 2B, left).

We compared J2-enriched peaks between mock and Aph+ATRi-treated cells and identified regions within the 5’ETS and internal spacer 1 (ITS1) of pre-rRNA as the most prominent dsRNAs induced by replication stress (Fig. 2, B and C). Consistent with J2’s specificity for long (>~40 bp) dsRNA, mature 18S and 5.8S rRNAs were depleted from the J2 pull-down despite harboring surface-exposed stem loops (Fig. 2C). Similarly, other 5’ppp-RNA induced by Aph+ATRi, such as RPPH1 (Fig. 1E), were not enriched in the J2 pull-down. In addition, other dsRNA species detected in J2 pull-down, including mitochondrial dsRNA or Alu dsRNAs, did not harbor 5’ppp/5’pp (Fig. 1E) and showed minimal changes upon Aph+ATRi (FC<2) (Fig. 2B).

To examine whether pre-rRNA duplexes can also activate MDA5, we performed previously established MDA5 filament footprint-seq assay^28^. In this assay, cytosolic RNA was incubated with recombinant MDA5, which selectively forms filament on dsRNA^1^, followed by extensive RNase A digestion, RNA size selection and sequencing to identify long stretches of dsRNA protected by MDA5 filament assembly (Fig. 2D, left). This analysis also identified 5’ETS and ITS1 of pre-rRNA as two most prominent RNA species protected by MDA5 filaments and enriched upon Aph+ATRi treatment (Fig. 2D, right). These results thus suggest aberrant rDNA-derived transcripts as potential ligands for both RIG-I and MDA5 during replication stress.

To further test the role of rDNA-derived transcripts in RLR activation, we used inhibitors for RNAP I (BMH21^33^), RNAP II (α-amanitin^34^) and RNAP III (ML60218^35^). Note that BMH21 acts through intercalating into rDNA^33^ and that no small molecule inhibitor is available to directly target RNAP I. BMH21 markedly reduced both sense and antisense 5’ETS RNA level in response to Aph+ATRi, as measured by RNA-FISH (Fig. 2E, and fig. S3, A and B). In contrast, neither α-amanitin nor ML60218 reduced 5’ETS RNA. Importantly, BMH21 reduced Aph+ATRi-mediated induction of *IFNB1, RANTES* mRNA without affecting housekeeping genes (e.g. *GAPDH*), whereas ML60218 had little effect (Fig. 2F). BMH21 also decreased pSTAT1 induction upon Aph+ATRi treatment (fig. S3C), further supporting the role of rDNA transcripts in RLR activation.

To further validate this conclusion, we examined the impact of depleting RPA194, the catalytic subunit of RNAP I. Since RPA194 is essential, we generated an RPA194-deficient RPE-1 cell line stably expressing degradable RPA194 by fusing with FKBP12^F36V^ (RNAP I-deg cells, Fig. 2G). Treatment with the degrader dTAGv-1^36^ selectively depleted RPA194-FKBP12^F36V^ in the RNAP I-deg cells, while showing little effect on endogenous RPA194 in the parental cells (Fig. 2G). Degradation of RPA194-FKBP12^F36V^ was accompanied by reduction in both sense and antisense 5’ETS RNA (fig. S3D) and RLR signaling, as measured by pSTAT1 level (Fig. 2G) or *IFNB1*/*RANTES* mRNA (fig. S3D). In the parental cells, dTAGv-1 reduced neither aberrant rRNA transcription nor RLR signaling (Fig. 2G, and fig. S3D).

Together, these results demonstrate that RLRs are activated by rDNA-derived transcripts during replication stress.

### ATR and CDK1 differentially regulate RLR signaling via rDNA transcription, independent of micronuclei or mitosis

Previous studies have shown that replication stress-induced innate immune activation is amplified by ATRi and suppressed by CDK1i, regardless of whether signaling is mediated by RLRs or cGAS^13,14^. Because ATR and CDK1 respectively promote and inhibit mitotic progression, it has been proposed that mitosis is required for innate immune activation^9,10,13^. Specifically, micronuclei formed when damaged chromosomes mis-segregate during mitosis, have been suggested as a source of cGAS^9,10^ and RLR activation^37^ under various genotoxic stress, although the relative contribution of micronuclei to innate immune signaling remains debated^38,39^.

We first confirmed the impact of ATRi and CDK1i on replication stress-induced RLR activation, as measured by *IFNB* and *RANTES* transcriptional induction or pSTAT1 level (Fig. 3A, and fig. S4A). The suppressive effect of CDK1i was recapitulated by siRNA-mediated depletion of CDK1 (fig. S4, B and C), excluding potential off-target effect of the inhibitor^40^. Moreover, CDK1i remained suppressive even when cells were synchronized in G1 and released into the S phase at which point CDK1i and Aph+ATRi were applied (fig. S4C), ruling out the possibility that CDK1i blocks immune activation simply by arresting cells in G2 before they can enter S phase under replication stress.

**Fig. 3.**
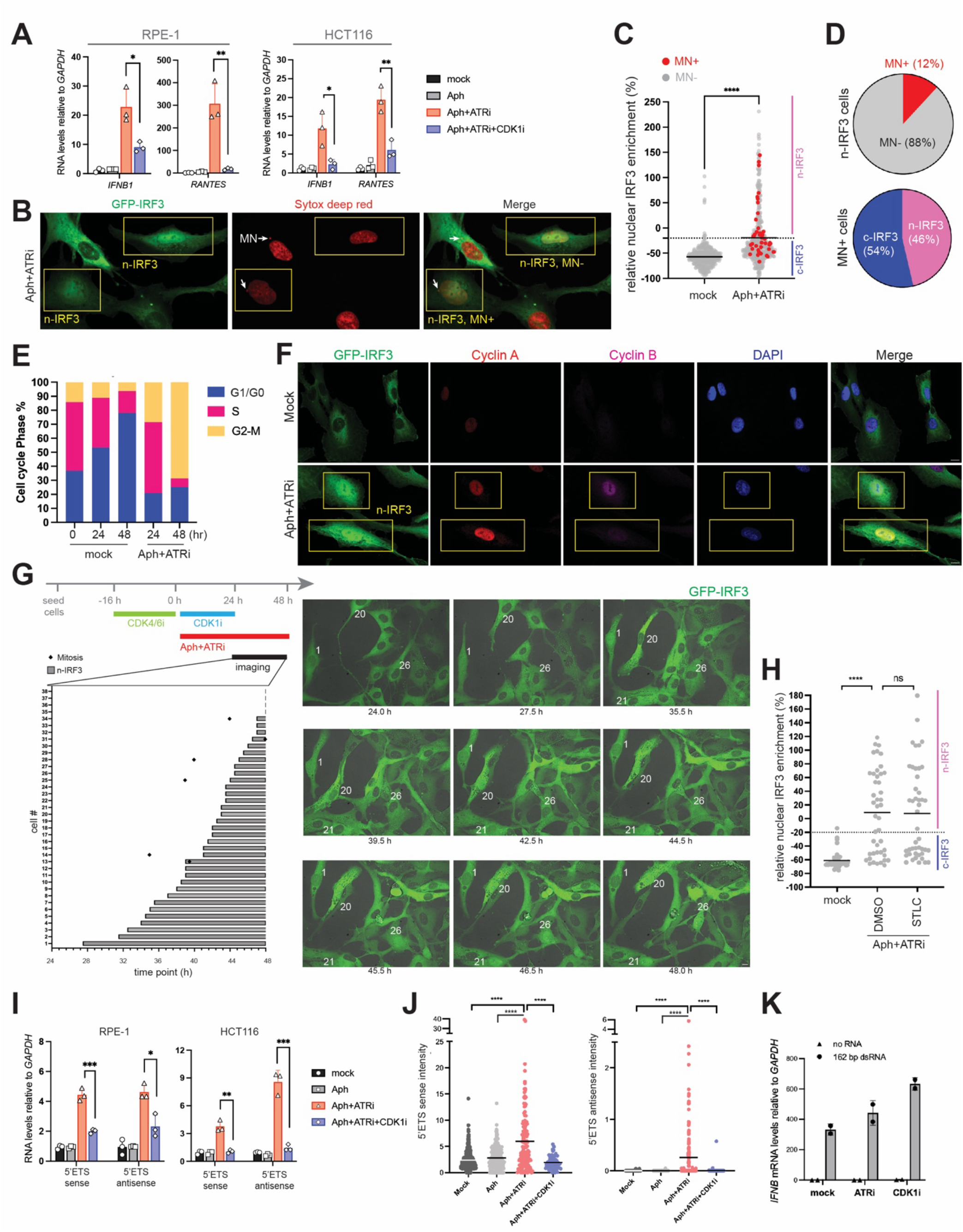
RLR activation correlates with aberrant rDNA transcription independent of micronuclei or mitosis. (A) The impact of ATRi (VE-821, 2.5 μM) and CDK1i (RO-3306, 6 μM) on antiviral signaling, as measured by RT-qPCR of *IFNB* and *RANTES* mRNAs relative to *GAPDH* in RPE-1 and HCT116 cells. Cells were treated with indicated drugs together with Aph (0.5 μM) for 48 h. Values represent mean ± SD from 3 biological repeats. \*\*\**p*<0.001, \*\**p*<0.01, \**p*<0.05 from two-tailed student’s t test. (B) Representative immunofluorescence images (Z-stacked) of RPE-1 cells treated with Aph+ATRi for 32 h. Sytox Deep Red stain indicates nuclei and micronuclei (MNs; white arrows). Cells with nuclear GFP-IRF3 (n-IRF3) are indicated with yellow boxes. (C) Quantification of nuclear IRF3 enrichment in mock and Aph+ATRi-treated cells. Nuclear enrichment (%) and n-IRF3 cells were defined as in Fig. 1C. Each point represents an individual cell (n=388 for mock, 497 for Aph+ATRi) from z-stacked images, where MN+ and MN-cells were colored red and gray, respectively. \*\*\*\**p*<0.0001 by two-tailed Mann–Whitney test. (D) Pie charts showing the proportion of MN+/-cells among nuclear IRF3 (n-IRF3) cells (top) and the proportion of n-IRF3 or c-IRF3 cells among MN+ cells (bottom). Both pie charts were derived from (C). (E) Cell cycle distribution of RPE-1 cells following Aph+ATRi treatment, determined by FxCycle Violet and Click-iT EdU staining. See fig. S5A. (F) Representative images of GFP-IRF3 (green), Cyclin A (red), Cyclin B (magenta), and DAPI (blue) in RPE-1 cells 32 h post Aph+ATRi or mock treatment. Yellow boxes highlight cells with n-IRF3. (G) Time-lapsed imaging of GFP-IRF3 translocation relative to mitosis. Top left: RPE-1 cells were synchronized at G1 with CDK4/6i (Palbociclib, 0.2 μM) and released into S phase in the presence of Aph+ATRi and CDK1i. CDK1i was added to block both G2>M transition and RLR activation. Upon CDK1i release, cells were subjected to Aph+ATRi for another 24 hr, during which images were acquired every 30 min. Bottom left: 34 cells showing IRF3 nuclear translocation were numbered in the order of translocation. Gray bars indicate duration of nuclear GFP-IRF3. ♦ marks mitosis (as measured by cell rounding). Right: representative time-lapsed images of GFP-IRF3 (green) with cell identities indicated. (H) Quantification of nuclear IRF3 enrichment (%) in mock or Aph+ATRi-treated cells, with and without mitotic arrest agent (S-Trityl-L-cysteine, STLC, 2 μM). Each point represents an individual cell (n=288 for mock and 134 for Aph+ATRi) from ~30 fields of view per condition. Images were analyzed as in (C). \*\*\*\**p* < 0.0001; ns, not significant, by two-tailed Mann–Whitney test. (I) Impact of ATRi and CDK1i on 5’ETS sense and antisense RNA induction. Cells were treated with indicated drugs for 48 h as in (A). Values represent mean ± SD from 3 biological repeats. \*\*\**p* <0.001, \*\**p* <0.01, \**p* <0.05 using two-tailed student’s t test. (J) RNA-FISH analysis of sense and antisense 5′ETS RNA in HCT116 cells under mock or 48 h treatment of indicated drugs. Each point represents an individual cell (n=555, 235, 135, 81, 291, 244, 164, 75 from left to right). ~20 fields of view per condition were analyzed. *\*\*\*\*p<0*.*0001* by two-tailed Mann–Whitney test. (K) Impact of ATRi (VE-821, 2.5 μM) and CDK1i (RO-3306, 6 μM) on antiviral signaling stimulated with 5’ppp-dsRNA (162 bp, 0.5 μg). RPE-1 cells were transfected with dsRNA and treated with indicated drugs 4 h post-transfection for 24 h, followed by RT-qPCR analysis of *IFNB* and *GAPDH* (internal control).

To assess whether micronuclei contribute to RLR signaling, we analyzed micronuclei formation together with IRF3 nuclear localization at the single-cell level. Cells were imaged 32 h after Aph+ATRi treatment, when IRF3 translocation was robust while nuclear morphology remained largely intact (Fig. 3B). Although Aph+ATRi increased micronuclei frequency (Fig. 3C), 88% of n-IRF3 cells lacked micronuclei (Fig. 3, C and D) and micronuclei-positive cells were about evenly split between n-IRF3 (54%) and c-IRF3 cells (46%) (Fig. 3, C and D). We also examined the possibility of micronuclei being the source of aberrant rDNA transcripts, but little to no 5’ETS RNA-FISH signal was detected within micronuclei (fig. S4E). These results argue against a role for micronuclei in RLR activation.

We next examined cell cycle distribution by flow cytometry using FxCycle violet staining and EdU label as a measure of DNA content and synthesis, respectively (fig. S5A). In the absence of Aph+ATRi, cells gradually shifted toward G1/G0 over 48 h, with fewer than 10% remaining in G2/M (Fig. 3E), consistent with quiescence in confluent cultures. In contrast, Aph+ATRi led to pronounced G2/M accumulation (~70–80% by 48 h) (Fig. 3E, and fig. S5A). Removal of Aph+ATRi during EdU labeling did not alter this distribution (fig. S5B). Although ATR inhibition promoted G2/M>G1 transition at 24 h, this effect was not observed at 48 h (fig. S5C), suggesting that accumulated damage enforces G2/M arrest even in the absence of ATR.

Consistent with these results, immunofluorescence analysis showed that Aph+ATRi enriched G2 markers, including Cyclin A, Cyclin B, and high Ki67, accompanied by reduced EdU incorporation (S phase) and Cyclin D levels (G1 phase) (fig. S6A). Few cells showed phosphorylated H3 (M phase) with or without Aph+ATRi (fig. S6A). In line with G2 arrest, RNA-seq showed marked suppression of histone gene expression upon Aph+ATRi, recapitulating G2 arrest by CDK1i (fig. S4D).

To understand how cell cycle state is related to RLR activation, we analyzed IRF3 nuclear localization together with cell cycle markers. Approximately 54% of cells with n-IRF3 displayed high levels of Cyclin A or B (Fig. 3F, and fig. S6A), indicating a G2 state. The remaining n-IRF3 cells, although lacking strong Cyclin A or B signal, exhibited elevated Ki67 levels and reduced EdU incorporation relative to cells with c-IRF3 cells, and with little to no p-H3 signal (fig. S6A). These results suggest that RLR activation may occur in G2/G2-like states prior to mitosis.

To directly test whether RLR signaling can indeed occur independent of mitosis, we performed live-cell imaging using cell rounding as a morphological indicator of mitotic entry. Cells were synchronized in G1 using a CDK4/6 inhibitor (palbociclib), released into S phase in the presence of Aph+ATRi together with the CDK1i to prevent mitotic entry and RLR activation. After release from CDK1i, cells were imaged for up to 24 h while maintaining Aph+ATRi (Fig. 3G). Continued presence of Aph+ATRi after CDK1i release was required for IRF3 activation (fig. S6B), indicating that ongoing replication stress in the presence of CDK1 activity drives RLR signaling. IRF3 nuclear translocation occurred 4-24 h after CDK1i release. Among cells showing IRF3 translocation (n=34), the majority (n=28) did not enter mitosis during imaging and only four cells underwent mitosis prior to IRF3 translocation (Fig. 3G). Additionally, the mitotic spindle inhibitor STLC^41^ did not prevent IRF3 nuclear localization (Fig. 3H, and fig. S6C) or pSTAT1 accumulation (fig. S6B), further supporting that RLR activation is independent of mitotic progression.

Given the importance of rDNA transcription in RLR activation (Fig. 1 and 2), we next asked whether ATR and CDK1 regulate RLR signaling by modulating aberrant rDNA transcription, either directly or indirectly. RT-qPCR analysis showed that 5′ETS sense and antisense transcripts were increased by ATRi and decreased by CDK1i in both RPE-1 and HCT116 cells (Fig. 3I). Similarly, RNA-FISH analysis also showed the effect of ATRi and CDK1i on 5’ETS RNA concordant with RT-qPCR analysis (Fig. 3J). Consistent with the notion that ATR and CDK1 impact RLRs through endogenous RNA ligand production rather than downstream signaling pathways, RLR activation in response to exogenous 162 bp 5’ppp-dsRNA was not suppressed by CDK1i and only modestly perturbed by ATRi (Fig. 3K).

Together, these data demonstrate that replication stress-induced RLR activation is divergently regulated by ATR and CDK1 via aberrant rDNA transcription, largely independent of micronuclei formation or mitotic progression.

### rDNA damage induces aberrant rDNA transcription and RLR signaling

To understand how replication stress induces aberrant rDNA transcription, we considered emerging evidence that DNA breaks, although often associated with local transcriptional repression^42–44^, can also induce transcription from nearby regions in a context-dependent manner (*e*.*g*. pre-existing transcriptional activity)^45–50^. Such damage-induced transcription often occurs in a divergent, bidirectional fashion^51,52^, which can generate antisense transcripts that initiate at or near the break site and overlap with upstream sense transcripts. We therefore hypothesized that replication-associated DNA breaks at rDNA loci drive aberrant sense and antisense transcription. Under this model, the impact of ATRi on rDNA transcription and RLR signaling can be explained by ATR’s role in stabilizing replication forks and limiting DNA damage^4,5^. Conversely, we asked whether CDK1 may promote DNA damage under replication stress. This hypothesis was in line with the RNA-seq results showing that CDK1i suppressed not only RLR-dependent antiviral genes, but also RLR-independent immediate early genes (IEGs) known to be induced by DNA damage^53,54^ (fig. S4, D and F).

To test the impact of ATR and CDK1 on DNA damage during replication stress, we performed immunofluorescence analysis of γH2AX, a histone variant that accumulates on damaged DNA. Aph+ATRi markedly increased the global level of γH2AX with concomitant increase in n-IRF3 compared to Aph alone. By contrast, addition of CDK1i to Aph+ATRi significantly reduced both the γH2AX levels and n-IRF3 cells (Fig. 4A). Similarly, immunoblot analysis of γH2AX and activated (phosphorylated) forms of ATM and DNA-PKcs, which function upstream of γH2AX alongside ATR, also showed the same trend (Fig. 4B, and fig. S7A). Furthermore, the effect of CDK1i on γH2AX was again recapitulated by siCDK1 (Fig. 4C), ruling out the possibility of inhibitor off-target effect. These findings suggest that ATR limits, while CDK1 amplifies, the DNA damage response during replication stress.

**Fig. 4.**
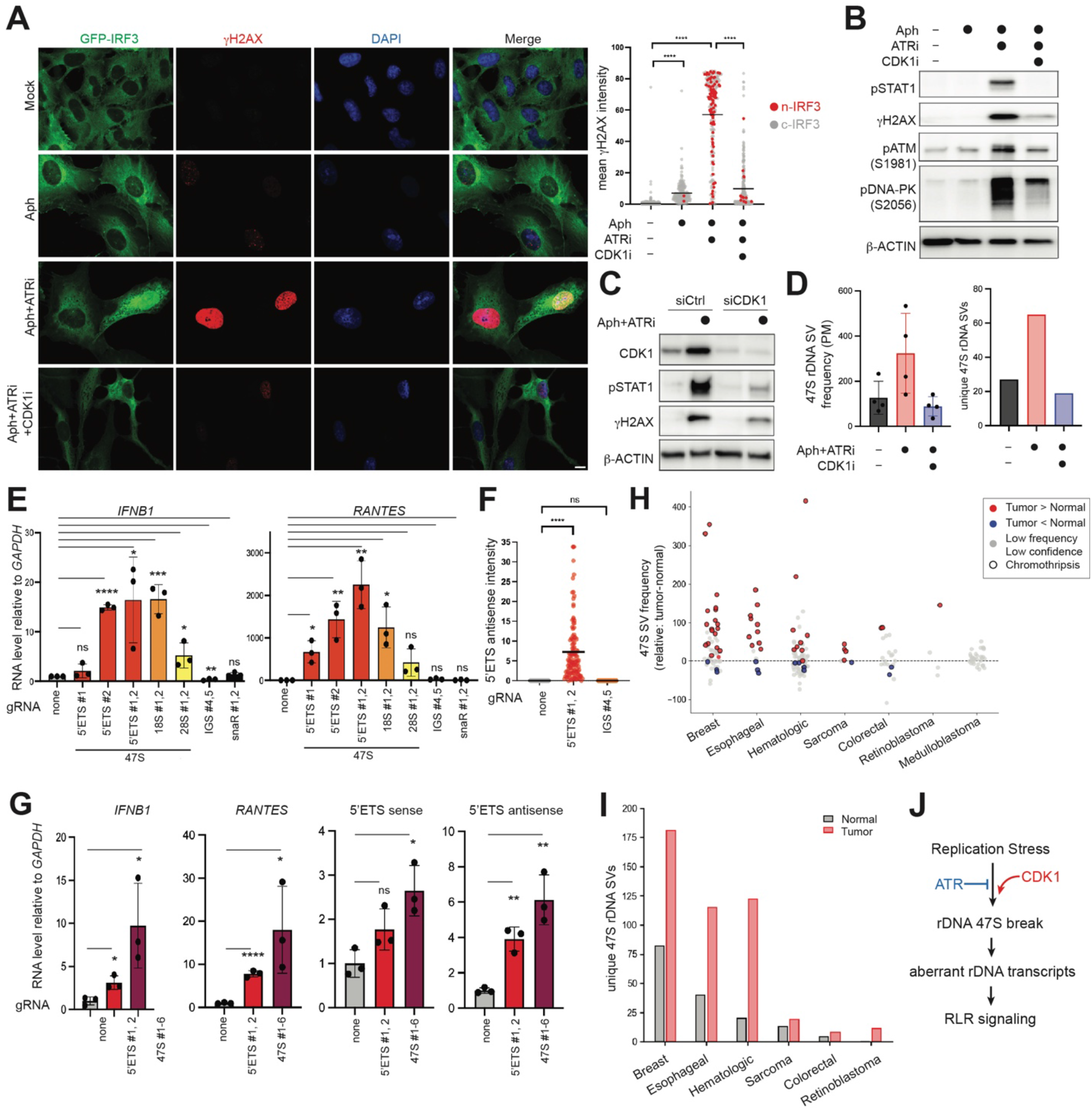
rDNA damage induces aberrant rDNA transcription and RLR signaling. (A) Representative images of GFP-IRF3 (green), γH2AX (red), and DAPI (blue) in mock, Aph, Aph+ATRi, and Aph+ATRi+CDK1i-treated RPE-1 cells (32 h). Right: Quantification of γH2AX intensity in cells with n-IRF3 (red) or c-IRF3 (gray). Each dot represents an individual cell (n=946, 209, 259 and 308 from left to right) from ~60 fields of view per condition. \*\*\*\**p* < 0.0001 by two-tailed Mann–Whitney test. (B) Western blot analysis of pSTAT1, γH2AX, pATM (S1981), and pDNA-PK (S2056) in RPE-1 cells treated with Aph, Aph+ATRi, and Aph+ATRi+CDK1i for 32 h. (C) Western blot analysis of pSTAT1, γH2AX, and CDK1 in RPE-1 cells transfected with control (siCtrl) or CDK1 siRNA (siCDK1), and treated with or without Aph+ATRi for 48 h. (D) Structural variations (SVs) in the 47S region of rDNA under replication stress with or without CDK1i. HCT116 cells were treated with indicated drugs for 48 h and whole genome sequencing (WGS) was performed. Sequences were mapped to rDNA repeat unit (IGS-47S), followed by SV analysis using Delly. All SV types (deletions, inversions and duplications) were aggregated, and only SV events overlapping with the 47S region were retained for downstream analyses. Left: 47S SV frequency as measured by reads supporting 47S SVs per million reads mapped to the 47S region (PM). Right: unique SV events in the 47S region. (E) Innate immune activation by Cas9-induced rDNA breaks. RPE-1 cells were electroporated with pre-assembled Cas9-gRNA complex targeting indicated regions in rDNA (5’ETS, 18S or 28S or IGS) or non-rDNA repeats (snaR), and harvested 48 h post-electroporation for RT-qPCR analysis. See fig. S7E and table S1 for gRNA target sites and sequences. \*\*\*\**p* <0.0001, ***p <0.001, \*\**p* <0.01, \**p* <0.05, not significant (ns) *p* > 0.05 by two-tailed student’s t test. (F) RNA-FISH analysis showing antisense 5′ETS in RPE-1 cells following Cas9-induced rDNA breaks 48 h. n = 367, 120, 186 cells for the no gRNA, 5′ETS #1,2, and IGS #4,5 groups, respectively, from ~20 z-stacked fields of view. *\*\*\*\*p* < 0.0001, not significant (ns) by two-tailed Mann–Whitney test. Scale bar, 10 µm. (G) Innate immune activation and rDNA transcription induced by Cas9-induced DNA breaks in HCT116 cells. Experiments were performed as in (E), with the exception of adding ATRi (2.5 μM) to cells immediately after electroporation. See fig. S7G for the comparison between with and without ATRi. \*\*\*\**p* <0.0001, \*\**p* <0.01, \**p* <0.05, not significant (ns) *p* > 0.05 by two-tailed student’s t test (H) Differential frequencies of SVs in 47S rDNA in tumor vs. matched normal samples. SV frequency was quantified as read counts supporting SVs per million reads mapped to 47S, which normalizes for variable rDNA copy numbers and different sequencing depths. Each data point represents an individual patient with paired tumor and normal whole-genome sequencing data and is colored red or blue depending on whether the tumor or normal sample shows higher SV frequency. Gray points indicate low-confidence events (<20 supporting reads or low frequency) in the sample with higher apparent SV levels. Patients highlighted in red or blue were further analyzed for chromothripsis, with black circle outlines indicating cases exhibiting chromothripsis. (I) Unique SV events in 47S region identified from patients showing a significant increase (red) in SV frequency in tumor vs matched normal samples from (H). For each cancer type, unique SV events from all such patients were aggregated. (J) Model of replication stress-induced RLR activation. Replication stress induces genomic instability, including at ribosomal DNA (rDNA) loci. rDNA breaks induce aberrant RNA polymerase I–dependent bidirectional transcription. The rDNA-derived sense and antisense transcripts form dsRNAs with 5’ppp that activate RLR signaling. ATR and CDK1 exert opposing effects on this pathway through their differential regulation of replication-associated DNA damage.

To examine whether DNA damage occurs at rDNA loci during replication stress, we analyzed previously published END-seq data^22^, which suggested enrichment of DNA breaks at the 47S region of rDNA relative to other highly transcribed housekeeping genes upon Aph+ATRi treatment (fig. S7B). Importantly, whole-genome sequencing showed that Aph+ATRi induced 47S rDNA structural variations (SVs), including inversions, deletions, and duplications consistent with end-joining repair, whereas CDK1i suppressed these events, whether looking at SV detection frequency or unique SV events (Fig. 4D). Similar effect of CDK1i was observed in both HCT116 (Fig. 4D) and RPE-1 (fig. S7C). Breakpoint analysis indicated that breaks are distributed across the 47S region without clear hot spots (fig. S7D). These SVs at 47S rDNA reflect 47S-specific DNA breaks, as demonstrated by Cas9-directed cleavage: targeting 47S, but not the non-transcribed intergenic sequence (IGS) within the rDNA repeat unit, induced SVs within 47S, not IGS, whereas targeting IGS induced SVs at IGS region, not 47S (fig. S8A). Thus, rDNA undergoes substantial DNA damage during replication stress in a CDK1-dependent manner, and that such damage can exceed the capacity of homologous recombination, leaving genomic scars in the form of SVs.

To determine whether rDNA damage alone is sufficient to induce aberrant rDNA transcription and RLR activation, we introduced targeted rDNA breaks using Cas9. Targeting one or two sites within the transcribed 47S pre-rRNA region, such as 5’ETS, robustly induced *IFNB* and *RANTES* expression (Fig. 4E) as well as production of 5’ETS antisense RNA (Fig. 4F) in RPE-1 cells. In contrast, targeting IGS within the same rDNA repeat failed to induce *IFNB* or *RANTES* expression (Fig. 4E) or 5’ETS antisense RNA production (Fig. 4F), despite equivalent γH2AX accumulation (fig. S8B). Targeting another clustered repetitive element (snaR)^55^ outside rDNA loci likewise failed to induce antiviral signaling (Fig. 4E). Consistent with a role for aberrant rDNA transcription in innate immune signaling, degradation of the RNAP I subunit RPA194 abrogated antiviral signaling induced by 5’ETS-gRNA (fig. S8C).

Similar innate immune activation was observed in HCT116 cells upon targeting the 47S rDNA region, accompanied by concordant increase in both sense and antisense 5’ETS RNA (Fig. 4G). Note that, unlike RPE-1, HCT116 cells required ATR inhibition for robust immune activation and 5’ETS RNA production (fig. S8D), which correlated with enhanced gH2AX signal upon ATR inhibition (fig. S8E), suggesting that Cas9-induced break triggers replication stress that is further amplified by ATRi. Consistent with a requirement for RLR signaling, MAVS knockout abrogated *IFNB* induction, while preserving aberrant 5’ETS RNA production (fig. S8F). Together, rDNA break is sufficient to trigger aberrant rDNA transcription and downstream RLR activation.

Finally, we asked whether 47S rDNA damage occurs under physiological conditions, including tumor development, as tumors are often subjected to high level of replication stress. Taking advantage of the fact that rDNA damage can leave genetic scars in the form of genomic SVs, we analyzed WGS data from 178 tumor–normal pairs^56–68^ across multiple tumor types. Although rDNA SVs were also detectable in normal samples, tumors generally showed increase in SV frequency and unique SV events within the 47S rDNA region, with particularly prominent changes in breast cancer, esophageal and hematologic cancers (Fig. 4, H and I, and fig. S8G). Nearly all tumors with elevated 47S rDNA instability also exhibited chromothripsis (Fig. 4H), a catastrophic form of genomic rearrangement that frequently drives tumor evolution. Notably, the magnitude of increase in both 47S SV detection frequency and unique SV events was comparable to that observed following Aph+ATRi treatment (Fig. 4D), supporting the physiological relevance of Aph+ATRi-induced rDNA damage.

Altogether, our results show that rDNA instability is sufficient to drive aberrant rDNA transcription and RLR-mediated inflammatory signaling, and is common in physiological genotoxic stress, such as tumor development.

## Discussion

Transcription and DNA damage repair are tightly interconnected. DNA damage has traditionally been thought to transiently suppress local transcription to avoid interference with repair processes^42–44^. However, accumulating evidence suggests that dysregulated transcription can be induced at damaged sites and may even facilitate repair in certain contexts^45–50^. This damage-induced transcription may arise from increased chromatin accessibility during repair^69^ or direct recruitment mechanisms^70^, and may be reconciled with damage-induced transcriptional suppression if canonical transcription is suppressed while aberrant transcription is induced at DNA lesions.

Here, we identify a previously unrecognized consequence of damage-induced transcription: innate immune signaling through aberrant RNA production from rDNA (Fig. 4J). Replication stress or double-strand breaks within rDNA leads to accumulation of otherwise unstable 5’ppp-containing pre-rRNA fragments together with overlapping antisense transcripts, which form dsRNA. A subset of these RNAs relocalize to the cytoplasm through an as-yet undefined mechanism, where they act as potent endogenous ligands for RIG-I and MDA5 and activate innate immune signaling.

Although damage-induced transcription can occur genome-wide, rDNA is uniquely poised as a potent source of immunostimulatory RNA under replication stress. Its high transcriptional capacity, multi-copy organization (~300-400 repeats) and nucleolar clustering likely facilitate both damage-induced transcription and interactions between sense and antisense RNA arising from distinct rDNA repeat units. Consistent with this, short dsRNAs processed by Dicer have been observed following endonuclease-induced DNA breaks at rDNA, but not at other sites^21^. In addition, rDNA is transcribed by RNAP I, and thus the transcripts retain 5′ppp ends required for RIG-I activation, whereas RNAP II transcripts are capped and largely inert. While RNAP III transcripts also carry 5′ppp ends, their dispersed genomic organization may limit efficient duplex formation, consistent with their limited recovery in our J2 pull-down assays. Finally, despite the largely co-directional organization of transcription and replication at rDNA loci^20^, rDNA remains susceptible to replication stress, as evidenced by frequent rearrangements in tumors and under experimental replication stress condition. While rDNA copy number variability has been documented in cancer, aging and replication stress^20,21^, our findings reveal a distinct functional consequence of rDNA instability in driving RLR signaling.

Our findings also revealed unexpected role of CDK1 in amplifying replication stress-associated DNA damage and the production of immunostimulatory RNA (Fig. 4J). In keeping with this, a recent report showed that CDK1 exacerbates replication defects at common fragile sites and contributes to their destabilization during replication stress^71^. These findings align with growing evidence that CDK1 functions extend beyond mitotic progression^72^, and can explain the widely observed phenomenon that CDK1 is required for both RLR and cGAS activation under genotoxic stress^13,14^. This also challenges a widely held view that mitotic progression is required for their activation, either by promoting micronuclei formation or by releasing nuclear content to cytosolic receptors^9,10^. It remains to be addressed how the aberrant rDNA transcripts access the cytoplasm and whether cells subjected to prolonged replication stress exhibit compromised nuclear integrity or altered nuclear export.

Altogether, our study identifies rDNA—a largely overlooked part of the genome—as a genomic element that converts cumulative replication-associated lesions into RNA-based innate immune signals, establishing a new link between genome integrity and innate immunity.

## Acknowledgements

We thank all members of the Hur lab for their helpful discussion and feedback. This study was supported by Howard Hughes Medical Institute (S.H.). Fluorescence imaging was performed at the Core for Imaging Technology & Education at Harvard Medical School, and flow cytometry samples were analyzed at the Boston Children’s Hospital FICR-PCMM Flow and Imaging Cytometry Core. Next-Gen Sequencing was done at Biopolymer Facility (Harvard Medical School).

## Author Contributions

IW, SA, SF performed experiments. XW performed bioinformatic analysis. IW and RC analyzed cellular images. SH supervised the project. IW, XW, SA and SH wrote the manuscript.

## Supplementary Materials

Figs. S1 to S8; Tables S1 to S5

## Materials and Methods

### Cell lines and culture conditions

HCT116 was grown and maintained in McCoy’s 5A (Modified) medium while RPE-1 cells were grown and maintained in DMEM medium, each supplemented with 10% fetal bovine serum (FBS). Wild type and Δp53 HCT116 and RPE-1 were kind gifts from Roger Greenberg lab (U. Penn). Cells were frequently tested for Mycoplasma contamination. The RPE-1 RNAP I-degron cells were generated by first overexpressing RPA194 with a C-terminal FKBP(F36V) under EF-1α promoter using lentiviral transduction, after which the endogenous *PolR1A* gene encoding RPA194 was knocked out using CRISPR-Cas9 (see below) using guide RNA sequences in table S1. The overexpressed RPA194 harbors a mutation in the PAM sequence, rendering it resistant to the editing. The RNAP I-degron cells were selected using 10 μg/ml Blasticidin and validated using Western blot analysis. All other knockout cell lines were also made by CRISPR-Cas9 (see below) using guide RNA sequences in table S1. Transient CDK1 knockdown was achieved by transfecting cells with siRNA (ON-TARGETplus human SMARTpool; Dharmacon, L-003224-00-0005; sequences listed in table S2) using Lipofectamine RNAiMAX (Thermo Fisher Scientific) according to the manufacturer’s instructions. An ON-TARGETplus non-targeting control siRNA (Dharmacon, D-001810-10-05) was used as a negative control. The siRNA sequences are provided in table S2.

### Plasmids

The mammalian RPA194 overexpression plasmid was generated by inserting the *PolR1A* gene in the pLenti-FKBP(F36V) plasmid using restriction-free cloning. The PAM sequence used for making PolR1A knock out was mutated using site-directed mutagenesis. pTRIP-GFP-IRF3 was a gift from Nicolas Manel (Addgene plasmid #127663). The DUSP11 constructs were kind gifts from Christopher Sullivan lab (UT Austin).

### RNAs

All RNAs used in this study were generated by *in vitro* T7 transcription as reported previously^73^. Templates for 47S 5’-ETS sense and antisense RNA were prepared by amplifying the first 241 nucleotides from the 47S pre-rRNA transcribed region. The RNAs were transcribed from the template using a mutant form of T7 RNAP that was reported to give low ssRNA and dsRNA by-products^74^. The RNA purity was assessed with TBE-PAGE analysis and the specific band was extracted from the gel using gel-crushing method followed by RNA clean-up using column (Zymo Research). For making sense:antisense hybrid, the two strands were pooled in equimolar ratio and heated to 95 °C followed by slowly cooling down to 25 °C at a ramp speed of−0.5 °C/s. The hybrid formation was confirmed by native TBE-PAGE. The positive control dsRNA sequences (162 bp and 512 bp) were derived from the first 150 and 500 bp sequences from human *IFIH1* and produced as previously reported^73^.

### Lentiviral production and transduction

For stable GFP-IRF3 expression, lentiviral particles were produced in 293T cells by co-transfecting psPAX2, pCMV-VSV-G, and pTRIP-GFP-IRF3 plasmids in 6-well plates using TransIT-293 Transfection Reagent (Mirus, MIR2700). Medium was replaced 24 h post-transfection with DMEM supplemented with 10% FBS. Viral supernatants were collected 24–48 h after medium change and filtered through 0.45 μm filters. Fresh viral supernatant was used to transduce RPE-1 Δp53 cells in the presence of polybrene (8 μg/ml). Medium was replaced with fresh growth medium after 6 h. At 72 h post-transduction, GFP-positive cells were isolated by fluorescence-activated cell sorting (FACS).

For inducible Flag–RIG-I expression, the Flag–RIG-I cDNA was subcloned into the doxycycline-inducible lentiviral vector pINDUCER20. Lentiviral particles were produced in 293T cells by transient transfection according to the manufacturer’s instructions. RPE-1 Δp53 GFP-IRF3 cells were transduced with lentivirus encoding Flag–RIG-I and selected with G418. Transgene expression was induced with doxycycline (500 ng/ml) for 24 hr prior to downstream assays.

For stable DUSP11 overexpression, Flag–DUSP11 or the catalytic mutant Flag–mDUSP11 (Cys152Ser) was cloned into a lentiviral expression vector (pLenti). Lentiviral particles were produced in 293T cells and used to transduce RPE-1 Δp53 ΔDUSP11 cells.

### CRISPR-Cas9 targeting

The guide RNA sequences were designed either manually or using the online tool https://www.idtdna.com/site/order/designtool/index/CRISPR_SEQUENCE from IDT. The guide RNA sequences have been provided in table S1. The ribonucleoprotein complex was first formed in vitro by incubating the guide RNA:tracrRNA hybrid with Alt-R™ S.p. HiFi Cas9 Nuclease V3 (IDT) at room temperature for 10 min. The RNP complex was then delivered into HCT116 or RPE-1 cells via electroporation using Neon™ NxT Electroporation System (Thermo Fisher Scientific). Briefly, 0.5 × 10^6^ cells were washed twice with PBS before resuspension in R buffer and mixed with RNP complex. The resuspended cells were electroporated at 1,530 V for 20 ms with one pulse (HCT116) or at 1,300 V for 20 ms with two pulses (RPE-1). The electroporated cells were immediately added to pre-warmed media in a 12-well plate and grown at 37 °C for 24-72 h depending on the experiment.

### RT-qPCR

Total RNAs were extracted using Directzol RNA miniprep kit (Zymo research) followed by DNase digestion for 1 h at 37 °C to eliminate DNA completely. Post-DNase treatment, RNA was further cleaned up using clean-up columns (Zymo Research). The cDNA was synthesized using Superscript IV (Applied Biosystems) according to the manufacture’s instruction. For 47S sense and antisense RNA, RT was performed with gene-specific reverse and forward primers respectively. Real-time PCR was performed using a set of gene specific primers, SYBR Green Master Mix (Applied Biosystems), and the CFX96 Real-Time PCR Systems (Biorad). For cytosolic sense/antisense 5’ETS RNA measurement, cells were grown and subjected to Aph+ATRi in a 15 cm dish for 48 h before lysis in hypotonic buffer (10 mM Tris pH 7.5, 10 mM KCl, 1 mM EDTA) using 50 strokes of Dounce homogenizer. The lysate was centrifuged successively at 1000g, 5000g, and 18,000g eliminating the pellet from each spin and recovering the final supernatant consisting of the cytosolic fraction. Trizol reagent was added to the samples and the RNA extracted by chloroform extraction followed by isopropanol precipitation. The purified RNA was treated with DNase I (NEB) for 1 h at 37 °C to eliminate DNA completely, and further purified by phenol:chloroform extraction and isopropanol precipitation, followed by RT-qPCR using the primer sequences shown in table S3.

### IRF3 dimerization assay

This assay was performed as described previously^28^. Briefly, HEK293T cells were homogenized in hypotonic buffer (10 mM Tris pH 7.5, 10 mM KCl, 0.5 mM EGTA, 1.5 mM MgCl_2_, 1 mM sodium orthovanadate, 1X mammalian ProteaseArrest, GBiosciences) and centrifuged at 1000 g for 5 min to pellet the nuclei. The supernatant (S1), containing the cytosolic and the mitochondrial fractions, was used for in vitro IRF3 dimerization assay. The stimulation mix for IRF3 activation was prepared by mixing 10 ng/μl RIG-I or MDA5, 2.5 ng/μl K63-Ub_n_ with 0-2.5 ng/μl RNA and pre-incubated at 4 °C for 30 min in 20 mM HEPES pH 7.4, 4 mM MgCl_2_ and 2 mM ATP. ^35^S-IRF3 was produced by in vitro translation using T7 Coupled Reticulocyte Lysate System (Promega) according to manufacturer’s instructions. The IRF3 activation reaction was carried out by adding 2 μl of stimulation mix to 20 μl reaction mixture containing 10 mg/ml of S1, 0.5 μl ^35^S-IRF3 in 20 mM HEPES pH 7.4, 4 mM MgCl_2_ and 2 mM ATP, and incubated at 30 °C for 1 h. Subsequently, the samples were centrifuged at 18,000 g for 5 min and the supernatant subjected to native PAGE analysis. IRF3 dimerization was visualized by autoradiography and phosphorimaging on Amersham Typhoon Imager (Cytiva).

### 5′ppp-RNA-seq

Total RNA from RPE-1Δp53 and HCT116Δp53 cells (mock or Aph+ATRi, 48h) was isolated using TRIzol (Invitrogen) followed by purification with Zymo-Spin II columns (Zymo Research). RNA was subjected to rRNA depletion using depletion beads (Thermo Fisher Scientific, A39115024) and further purified using Zymo-Spin IC columns. To enrich for 5′-triphosphorylated RNA species, samples were heat at 95 °C for 5 min and treated with XRN1 (New England Biolabs, M0338L) at 37 °C for 2 h in the presence of RNase inhibitor, followed by TRIzol cleanup. RNA 5′ ends were converted using RNA 5′ polyphosphatase (Rpp; Lucigen, RP8092H) at 37 °C for 1 h. RNA was fragmented by heat at 94 °C for 4 min; New England Biolabs, E6150S) and purified, followed by 3′ end dephosphorylation using T4 polynucleotide kinase (Epicentre, P0503K) at 37 °C for 30 min. Libraries were prepared using the Collibri™ Stranded RNA Library Prep Kit (Thermo Fisher Scientific) according to the manufacturer’s instructions, including adapter hybridization, ligation, reverse transcription, and PCR amplification. Final libraries were purified and subjected to Illumina paired-end sequencing (PE150).

For Oxford Nanopore sequencing, the same 5′ppp RNA-seq workflow was followed, except that RNA fragmentation was omitted. Following capture with the Collibri™ Stranded RNA Library Prep Kit, 5′ppp-containing 5′ETS fragments were enriched by 10 cycles of PCR using a TruSeq Read 1–5′ETS fusion forward primer (CTCTTCCGATCT<u>GCTGACACGC</u>) and an i7 reverse primer (CAAGCAGAAGACGGCATACGAG). Amplified DNA (200 fmol) was end-repaired and dA-tailed (NEBNext Ultra II, E7546; 20 °C 5 min, 65 °C 5 min), purified with AMPure XP beads, and quantified by Qubit dsDNA HS assay. Native barcodes (SQK-NBD114.24) were ligated (NEB M0367, 25 °C, 20 min), pooled, and purified (0.8× AMPure XP). Sequencing adapters were ligated (NEB E6056, 25 °C, 20 min), and libraries were loaded onto a MinION R10.4.1 flow cell (Oxford Nanopore Technologies).

### J2 pull-down RNA-seq

RPE-1Δp53 cells were grown 2 x 15 cm dish and subjected to Aph+ATRi treatment for 48 h. The cells were washed twice with ice-cold PBS and lysed in the buffer (50 mM Tris pH 7.5, 100 mM NaCl, 3 mM MgCl_2_, 0.5% NP-40) by incubating on ice for 5 min. Spin the lysate at 13,000g for 5 min and collect the supernatant. Set 50 μl aside for Input sample and add 10 μl mouse J2 antibody (1 mg/ml) and mix well. Add 50 μl pre-equilibrated Dynabeads Protein G magnetic beads (Invitrogen) and incubate at 4 °C for 3 h with constant gentle agitation. Wash the beads thrice with lysis buffer and then incubate with RNase I_f_ for 15 min at RT before adding Trizol to the beads and subjecting to RNA extraction with Direct-zol RNA miniprep kit (Zymo Research). The purified RNA was used for cDNA library preparation using SMARTer Stranded Total RNA-Seq Kit v3 - Pico Input Mammalian according to the manufacturer’s instructions. The cDNA library was sequenced using the Illumina Partiallane NovaSeq or NextSeq platform with paired-end 150 bp reads.

### MDA5 filament footprint-seq

The assay was performed as previously reported^28^. Cytosolic RNA (5 ng/μl) purified from RPE-1 Δp53 cells (mock or Aph+ATRi) was pre-incubated with MDA5Δ2CARD (150 nM) at RT for 10 min in 20 mM HEPES pH 7.5, 50 mM NaCl, 2 mM MgCl_2_ and 2 mM DTT followed by addition of RNase A (0 or 0.5 ng/μl) and incubation for another 5 min at RT. The RNase A digestion was quenched by adding three volumes of TRIzol reagent (Thermo Fisher) and the RNA was purified with Directzol RNA miniprep kit (Zymo Research) using manufacturer’s protocol. The extracted RNA was further purified using QIAquick PCR purification kit (QIAGEN) to remove small digestion products and the purified RNA was used for the cDNA library preparation using SMARTer Stranded Total RNA-Seq Kit v3 - Pico Input Mammalian according to the manufacturer’s instructions. The cDNA library was sequenced using the Illumina Partiallane NovaSeq or NextSeq platform with paired-end 150 bp reads.

### Total RNA-seq

HCT116Δp53 were grown in a 12-well plate and synchronized in G1 with CDK4/6 inhibitor (Palbociclib; 0.2 μM) for 16 h. Cells were then washed twice with PBS followed by mock or Aph+ATRi treatment for 48 h in the presence or absence of CDK1 inhibitor (RO3306; 6 μM). Total RNAs were extracted using Directzol RNA miniprep kit (Zymo research) followed by DNase digestion for 1 h at 37 °C to eliminate DNA completely. RNA was then further purified using clean-up columns (Zymo Research) and was used for cDNA library preparation with SMARTer Stranded Total RNA-Seq Kit v3 - Pico Input Mammalian according to the manufacturer’s instructions. The cDNA library was sequenced using the Illumina Partiallane NovaSeq or NextSeq platform with paired-end 150 bp reads.

### Whole Genome Sequencing (WGS)

Genomic DNA was extracted using Monarch HMW DNA Extraction Kit for Cells & Blood (New England Biolabs). Extracted DNA was fragmented using NEBNext dsDNA Fragmentase (New England Biolabs) at 37 °C for 40 min. The fragmented DNA was purified using SPRI Select beads (Beckman) and DNA libraries were prepared using NEBNext Ultra II DNA library Prep Kit for Illumina (New England Biolabs) according to the manufacturer’s protocol. The library quality was assessed using TapeStation analysis (Agilent) and the concentration estimated by Qubit Fluorometer (Agilent). Deep sequencing was performed using a NextSeq or NovaSeq sequencer (Illumina) with paired-end 150 bp reads.

### Bioinformatics analysis

#### Total RNA-seq and differential gene expression analysis

Pair-ended reads were trimmed by trimmomatic^75^, and mapped to Grch38.p14 by STAR in splice-aware mode^76^. Resultant bam files were sorted with samtools^77^ by read coordinates. Sorted bam files were processed by Rsubread^78^, and reads were counted for human gene transcripts by Gencode V43^79^. Multi-mapping reads were counted by fraction. Read count files were processed in Deseq2^80^, and TPM values were used to calculate z-scores.

#### 5’ppp-RNA-seq, MDA5 footprint-seq, and J2 PD-seq analysis

Pair-ended reads were trimmed by trimmomatic^75^, and mapped to rDNA repeat unit (including 47S pre-rRNA and IGS) and then to repbase^81^ by Bowtie2^82^. The reads unmappable to rDNA/repbase were finally mapped to Grch38.p14^76^. Only primary alignments were kept by samtools^77^. To adjust for different sequencing depths among samples, the bam files were first converted to bedgraphs by bedtools genomecov^83^, and the bedgraph coverage values were normalized to per-million-mapped reads (excluding rDNA) by multiplying the coverage value with a factor of 10^6^/(total non-rDNA mapped read counts).

Normalized per-million (PM) bedgraph tracks between experiment and control group were subtracted by MACS3^84^ bdgcmp, and peaks were called based on subtracted PM bedgraph by MACS3 bdgpeakcall (-l 30, -c 20). Specifically, the -Rpp track was subtracted from the +Rpp track for 5’ppp-RNA-seq; -RNase A was subtracted from +RNase A for MDA5 footprint-seq; input was subtracted from pulldown track for J2 PD-seq. Peaks were annotated by overlapping with Hg38 gene annotations, as well as Repbase annotation for repeat regions. AuC of individual PM bedgraphs within peak regions, both experiment and control groups, were calculated by bedtools intersect and adding all bedgraph coverage values within peak regions.

#### WGS mapping and SV calling

Paired-end WGS fastq files were mapped to rDNA repeat unit (44 kb, IGS-47S) or Hg38 by bowtie2, and primarily mappable reads within IGS, 47S in rDNA, or in all of Hg38, were kept for normalization purposes. Position-sorted bam files were used as input for Delly SV discovery^85^, including DEL (deletion), DUP (duplication), INV (inversion) and TRA (translocation) events. Because TRA detection requires multiple reference sequences, it was performed only for hg38-based analyses. Only SVs with “PASS” quality in Delly vcf files were used for downstream analyses. Given the repeat nature of rDNA, DUP events longer than 44kb were excluded as artifacts due to mapping all rDNA reads onto a single consensus reference. For rDNA analysis, SVs that overlap with IGS or 47S region were separated, and the total reads supporting SV discovery were normalized to per-million reads primarily mapped to IGS or 47S.

#### Copy number and chromothripsis

Paired bam files of tumor and normal from the same patient ID were analyzed by CNVkit, and copy number were calculated by patient over normal^86^. Both the copy number and tumor vcf file called by Delly were used as input for Shatterseek^87^, and chromosomes with copy number oscillation >=7 with SV counts in cluster >6 were classified as positive for chromothripsis^87^.

#### dbGaP controlled data access and analysis

WGS of cancer patients and matched normal were downloaded from NIH dbGaP under project #42869: Innate Immune Activation during Genotoxic Stress, with 13 cohorts: phs000245.v1.p1^56^, phs000369.v1.p1^57^, phs000579.va.p1^58^, phs000472.v2.p1.c1^59^, phs000374.v1.p1^60^, phs000598.v2.p2^61^, phs000348.v2.p1^62^, phs000414.v1.p1^63^, phs000804.v2.p1^64^, phs000341.v2.p1^65^, phs000409.v1.p1^66^, phs000352.v1.p1^67^, and phs000340.v3.p1^68^.

### Immunoblotting

Cells were lysed in SDS lysis buffer (2% SDS, 50 mM Tris-HCl, pH 7.5, 150 mM NaCl, 10 mM DTT) supplemented with 1× Mammalian ProteaseArrest™ Protease Inhibitor Cocktail (G Biosciences, 786-433) and heated at 95 °C for 5 min. Protein concentrations of cleared lysates were determined using a BCA protein assay (Thermo Fisher Scientific). Equal amounts of total protein were mixed with 1× SDS sample buffer (62.5 mM Tris-HCl, pH 6.8, 10% glycerol, 2% SDS, 2.5% β-mercaptoethanol), boiled for 5 min, and resolved on 4–15% Tris–glycine gels (Bio-Rad, 4561086). Proteins were transferred onto 0.45 μm PVDF membranes (Thermo Fisher Scientific, 88518). Membranes were blocked in 5% (w/v) nonfat dry milk or 5% BSA in TBS containing 0.1% Tween-20 (TBST) for 1 h at room temperature. Blots were incubated with primary antibodies diluted in TBST overnight at 4 °C with gentle agitation, followed by incubation with HRP-conjugated secondary antibodies for 1 h at room temperature. Signals were detected using a ChemiDoc MP Imaging System (Bio-Rad).

The following primary antibodies were used: phospho-Histone H2A.X (Ser139) (Cell Signaling Technology, 9718), DUSP11 (Proteintech, 10204-2-AP), RPA194 (Santa Cruz Biotechnology, sc-48385), RIG-I (Cell Signaling Technology, 3743), MDA5 (Cell Signaling Technology, 5321), STING (Cell Signaling Technology, 13647), MAVS (Bethyl Laboratories, A300-782A), phospho-STAT1 (Cell Signaling Technology, 9167), STAT1 (Cell Signaling Technology, 14995), β-actin (Cell Signaling Technology, 8457), vinculin (Cell Signaling Technology, 4650), ISG15 (Cell Signaling Technology, 2758), CDK1 (Cell Signaling Technology, 9116), phospho-ATM (Ser1981) (Cell Signaling Technology, 13050), and phospho-DNA-PKcs (Ser2056) (Cell Signaling Technology, 68716).

### Cell cycle analysis by flow cytometry

Cell cycle analysis was performed using the Click-iT™ Plus EdU Alexa Fluor™ 488 Kit (Thermo Fisher Scientific, C10632) and FxCycle™ Violet Stain (Thermo Fisher Scientific, F10347) according to the manufacturer’s instructions. EdU was added to culture medium at a final concentration of 10 μM for 30 min prior to harvest. At the indicated time points, cells were washed once with PBS containing 1% BSA and pelleted at 300 × g for 5 min. Cells were fixed in 4% paraformaldehyde for 15 min at room temperature, permeabilized using 1× Click-iT™ permeabilization buffer for 15 min, and subjected to the Click-iT™ Plus reaction for 30 min at room temperature. DNA was stained with FxCycle™ Violet for 30 min. Samples were analyzed using a FACSCanto II flow cytometer (BD Biosciences) at the BCH FICR-PCMM Flow and Imaging Cytometry Core, and data were processed using FlowJo v10 software.

### Immunofluorescence

Cells were seeded on coverslips and treated as indicated. Cells were fixed in 4% paraformaldehyde for 15 min at room temperature and permeabilized with 0.5% Triton X-100 for 8 min. After blocking in PBS containing 5% normal serum and 0.1% Triton X-100 for 1 h at room temperature, cells were incubated with primary antibodies diluted in PBS containing 1% BSA and 0.1% Triton X-100 overnight at 4 °C. Following washes, cells were incubated with appropriate secondary antibodies for 1 h at room temperature. Nuclei were stained with DAPI. Coverslips were mounted using Fluoromount-G (SouthernBiotech, 0100-01) and imaged on a Nikon Ti motorized inverted microscope equipped with a Yokogawa CSU-X1 spinning disk confocal system using 40× or 100× oil-immersion objectives (Harvard Medical School Core for Imaging Technology & Education). Images were processed using Fiji and analyzed in Python.

Primary antibodies used were phospho-Histone H2A.X Ser139 (Cell Signaling Technology, 9718), Ki67 (Cell Signaling Technology, 9129), Cyclin A2 (Cell Signaling Technology, 67955), Cyclin B1 (Cell Signaling Technology, 12231), Cyclin D (Cell Signaling Technology, 55506), phospho-Histone H3 Ser10 (Cell Signaling Technology, 9706S), and fibrillarin (Thermo Fisher Scientific, MA3-16771). 5EU RNA labeling was performed using the Click-iT™ RNA Alexa Fluor™ 594 Imaging Kit (Thermo Fisher Scientific, C10330). SYTOX™ Deep Red Nucleic Acid Stain (Thermo Fisher Scientific, D1306) was used where indicated.

### Live-cell imaging

IRF3–GFP RPE-1 Δp53 cells were seeded in 12-well glass-bottom plates (Cellvis, P12-1.5H-N) and imaged under the indicated conditions using a Nikon Ti inverted fluorescence microscope equipped with a Yokogawa spinning disk confocal system and an incubation enclosure (Harvard Medical School Core for Imaging Technology & Education), using a 40× air objective. Cells were monitored over time as indicated.

### RNA Fluorescence In Situ Hybridization (RNA-FISH)

Sense 47S 5′ETS RNA (targeting TSS +1 to +242) was detected by single-molecule RNA FISH (Stellaris) in HCT116 and RPE-1 cells, whereas antisense 47S RNA (targeting TSS −1080 to +241) was detected by single-molecule RNA FISH in HCT116 cells and by signal amplification RNA FISH (ACD RNAscope) in RPE-1 cells. For Sterllaris RNA-FISH probes (table S4), oligonucleotides (IDT) were labeled using terminal deoxynucleotidyl transferase (TdT) with aminoallyl-dUTP at 37 °C overnight, purified by Zymo column cleanup, conjugated to Cy5-NHS in sodium bicarbonate buffer, and desalted using Bio-Rad P-4 beads. For Stellaris RNA FISH, cells were fixed in 4% paraformaldehyde for 10 min, permeabilized in 70% ethanol at 4 °C for 1 h, and hybridized overnight at 37 °C with pooled Stellaris probes. After washing, nuclei were stained with DAPI and samples were mounted in VECTASHIELD (VWR,101098-042). For RNAscope RNA-FISH, antisense RNA was detected using 12 pairs of adjacent “Z” probes with proprietary sequences targeting the TSS −1080 to +241 region. Cells were fixed in 4% paraformaldehyde for 30 min and dehydrated through an ethanol series (50%, 70%, 100%) followed by rehydration (100%, 70%, 50%). Endogenous peroxidase activity was quenched using RNAscope Hydrogen Peroxide for RT 10min. Cells were treated with diluted RNAscope Protease III (1:5 in PBS), followed by hybridization with custom probes at 40 °C for 2h. Signal amplification was performed using AMP1, AMP2, and AMP3 reagents, followed by HRP-C1 development and labeled with Opal 690 (Akoya Biosciences, FP1497001KT). After HRP blocker incubation, nuclei were stained with DAPI, and samples were mounted using ProLong™ Diamond Antifade Mountant (Thermo Fisher Scientific, P36965). Z-stacks were acquired using a Nikon Ti motorized inverted microscope equipped with a Yokogawa CSU-X1 spinning disk confocal system and a 40× oil-immersion objective (Harvard Medical School Core for Imaging Technology & Education).

### Image analysis

#### Preprocessing

Fluorescence microscopy images were acquired in .nd2 format and processed in ImageJ2 (v.2.14.0/1.54f) to generate single- and merged-channel TIFF files under identical acquisition settings, with brightness and contrast applied uniformly across conditions. For z-stacks, mean intensity projections were generated prior to analysis. TIFF images were converted to NumPy arrays (.npy) using *tifffile* (v.2025.9.9) and NumPy (v.2.2.6)^88^, then converted to grayscale; channels of interest were extracted, consistently ordered, and saved as standardized (C, H, W) arrays for downstream analysis.

#### Image analysis 1

For each condition, nuclei were segmented from the DAPI channel following Gaussian blurring and Otsu thresholding^89^ using OpenCV (v4.12.0). To define an appropriate size cutoff, nuclear areas were first measured from all segmented contours, and their global distribution was visualized as a histogram. Based on this distribution, nuclei with areas below 1500 px^2^ were excluded. To further eliminate irregular objects and debris, nuclei with circularity < 0.65^90,91^ were removed. For each retained nucleus, we calculated area, mean intensity, and total intensity for DAPI, anti-47S channels.

#### Image analysis 2

For each condition, nuclei were segmented from the DAPI channel following Gaussian blurring and Otsu thresholding using OpenCV (v4.12.0). Following the same process as Image analysis 1, only contours with areas > 3500 px^2^ and circularity > 0.65 were retained to ensure well-defined nuclear regions. For each nucleus, IRF3 mean intensity was measured inside the nuclear mask and compared to a cytoplasmic “ring” region created by dilating the nuclear contour outward (~5–15 px). Nuclear enrichment of IRF3 was quantified as the difference between mean nuclear and mean cytoplasmic intensities (Δ = nuclear – cytoplasmic). Cells were classified as nuclear-positive if nuclear IRF3 intensity exceeded cytoplasmic intensity, or if nuclear intensity was only slightly lower (within 20% of cytoplasmic intensity) to account for minor boundary misalignments.

#### Image analysis 3

For each condition, nuclei were segmented from the Sytox channel following Gaussian blurring and Otsu thresholding using OpenCV (v4.12.0). Following the same process as Image analysis 1, only contours with areas > 3500 px^2^ and circularity > 0.65 were retained to ensure well-defined nuclear regions. Each nucleus was refined by ROI-based contour matching on the Sytox channel, and micronuclei (MN) were identified from the Sytox channel within a surrounding ring (~100 px expansion) based on size (25–1000 px^2^) and circularity (> 0.65) criteria. Each retained nucleus was further categorized as micronucleus-positive (MN^+^) or micronucleus-negative (MN^−^). IRF3 mean intensity distributions were computed across all cells and stratified by micronucleus status (MN^+^ vs. MN^−^).

#### Image analysis 4

For each condition, nuclei were segmented from the DAPI channel following Gaussian blurring and Otsu thresholding using OpenCV (v4.12.0). Following the same process as Image analysis 1, only contours with areas > 3500 px^2^ and circularity > 0.65 were retained to ensure well-defined nuclear regions. Cells were classified as nuclear-positive if nuclear IRF3 intensity exceeded cytoplasmic intensity (or was only slightly lower, within 20%, as in Image analysis 3). To assess cell cycle activity, ki67 signal intensities were quantified per nucleus. Densities (total intensity per area) were computed for nuclear-positive and nuclear-negative cells, and distributions were compared across conditions. Other markers (cyclin B, cyclin A, cyclin D, phospho-histone H3, EdU, and γH2AX) were analyzed using the same pipeline. Cyclin A–high cells were defined as cells with nuclear cyclin A intensity exceeding the 80th percentile of the mock-treated control. In GS-treated n-IRF3 cells, 21 out of 39 cells (54%) met this criterion.

---

**Fig. S1.**
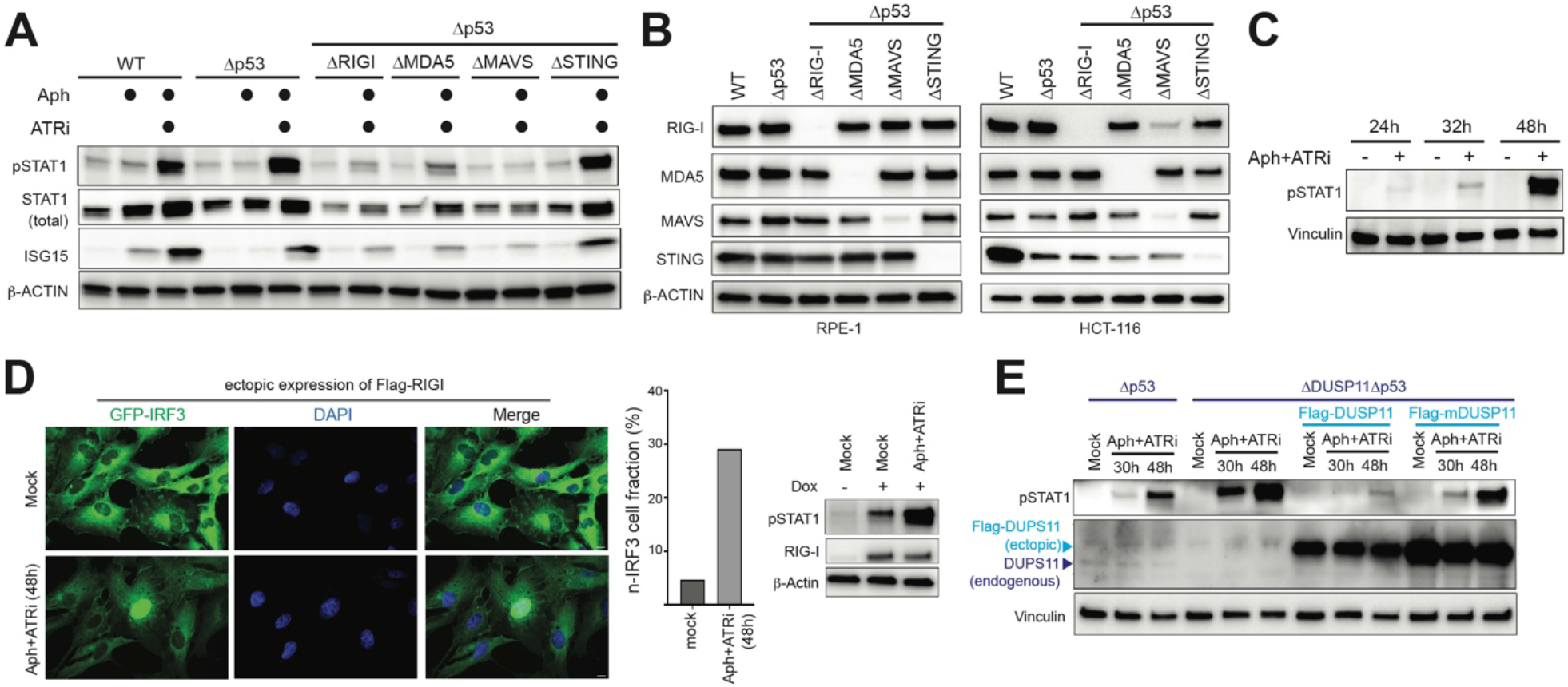
Replication stress activates RLRs over 32-48 h. (A) Western blot analysis of STAT1 phosphorylation (pSTAT1) and interferon-stimulated genes (e.g. ISG56, and ISG15) (right). HCT116 cells of indicated genotypes were treated with aphidicolin (Aph, 0.5 μM) and ATR inhibitor (VE-821; ATRi, 2.5 μM) for 48 h prior to analysis. β-ACTIN was used as a loading control. (B) Western blot analysis validating indicated genotypes in RPE-1 (left) and HCT116 (right) cells. Cells were treated with IFNb for 24 h to upregulate RIG-I and MDA5 that are otherwise too low to detect. (C) Western blot analysis of pSTAT1 in RPE-1 cells treated with Aph+ATRi for 24 h, 32 h and 48 h. (D) Representative immunofluorescence of GFP-IRF3 (green) and nuclei (DAPI, blue) in cells ectopically expressing Flag-RIG-I under mock or Aph+ATRi 48 h treatment. Bar plot shows percentage of cells with nuclear IRF3 (n-IRF3), as defined in Fig. 1C, from analysis of 148 and 120 cells for mock and Aph+ATRi treatment conditions, respectively. Immunoblot confirms RIG-I expression and pSTAT1 activation. (E) Western blot analysis of pSTAT1 in indicated RPE-1 cells with or without Flag-DUSP11 or catalytic mutant Flag-mDUSP11 (Cys152Ser) overexpression under mock or Aph+ATRi (30, 48 h) conditions. Vinculin, loading control.

**Fig. S2.**
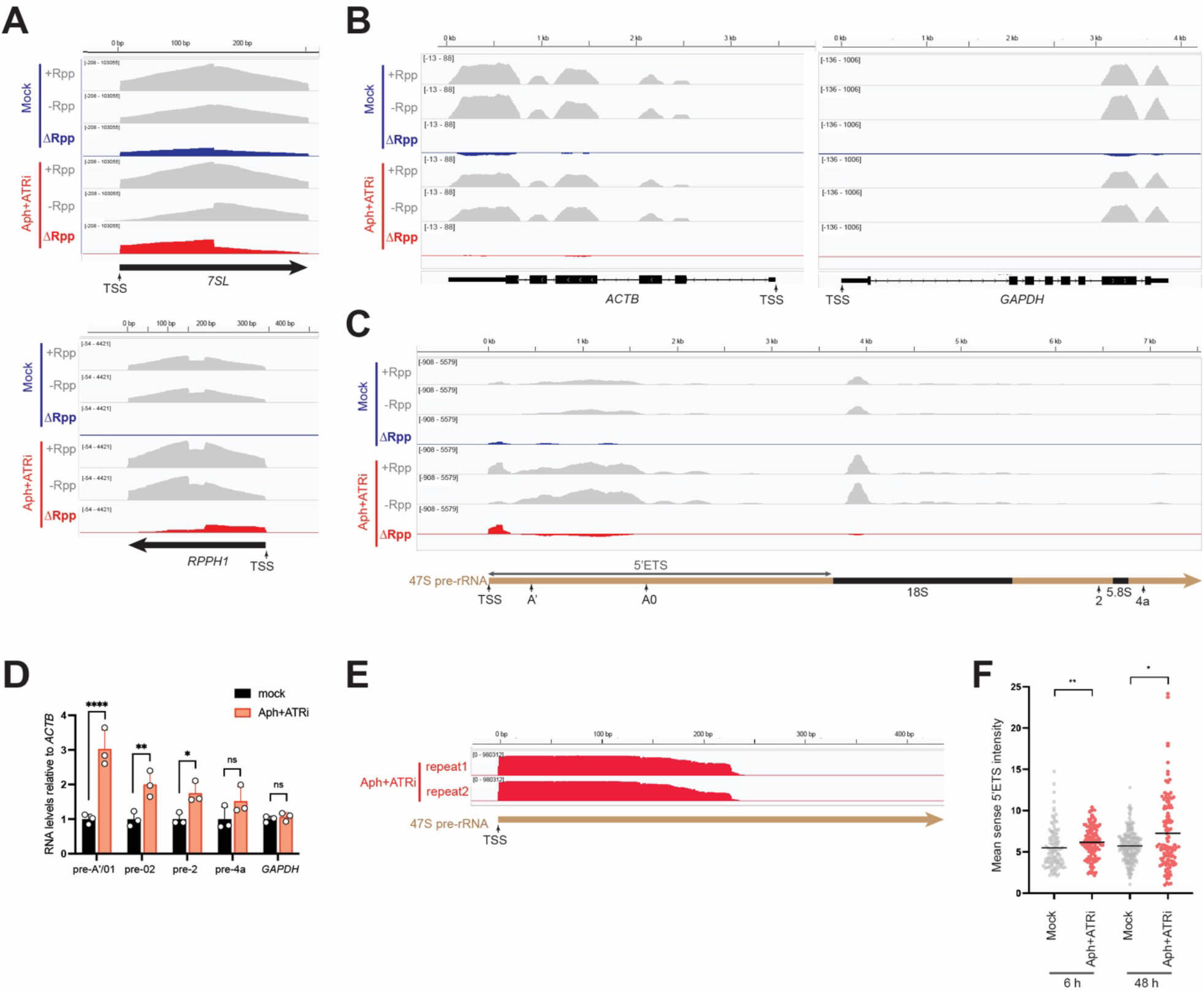
Replication stress induces dysregulated rDNA transcription and processing. (A-C). Representative IGV tracks of 5′pppRNA-seq. (A) 7SL (top) and RPPH1 (bottom), (B) *ACTB* (left) and *GAPDH* (right) in RPE-1 cells, and (C) the 47S pre-rRNA 5′ETS region in HCT116 cells. 5’ppp-RNAs were identified as Rpp-enriched regions by subtracting –Rpp from +Rpp coverage (ΔRpp), followed by peak calling on the ΔRpp track. See Fig. 1D for the workflow and Fig. 1E for the systematic analysis results in both RPE-1 and HCT116 cells. (D) RT-qPCR analysis showing levels of unprocessed 47S pre-RNA relative to *ACTB* in mock vs. Aph+ATRi-treated RPE-1 cells. PCR was performed using primers flanking indicated processing sites. All processing sites are shown in (C) except pre-02, which is at the junction between 28S rRNA and 3’ETS. See table S3 for primer sequences. Values represent mean ± SD from 3 independent experiments. *P*-values were based on two-tailed student’s t test. **** *p* = <0.0001, ** *p* = <0.01, * *p* = <0.05, not significant (ns) *p* > 0.05 (E) Nanopore sequencing of RNA harboring 5’ppp and 5’-end sequence of 47S pre-rRNA. 5’pppRNA-seq libraries were generated as in Fig. 1D without RNA fragmentation. 47S pre-rRNA 5’end sequences were then selectively PCR-amplified, during which nanopore sequencing adaptors were added. See Methods. Tracks from two replicates (repeat 1 and repeat 2) in Aph+ATRi-treated cells show read coverage that starts sharply at the known TSS and extending to ~241 nt downstream. (F) RNA-FISH analysis of sense 5′ETS RNA in RPE-1 cells under mock or Aph+ATRi treatment for 6 h and 48 h. Each point represents an individual cell (n=107,121,196, 122 from left to right). \**p*<0.05, \*\**p*<0.01 by two-tailed Mann–Whitney test.

**Fig. S3.**
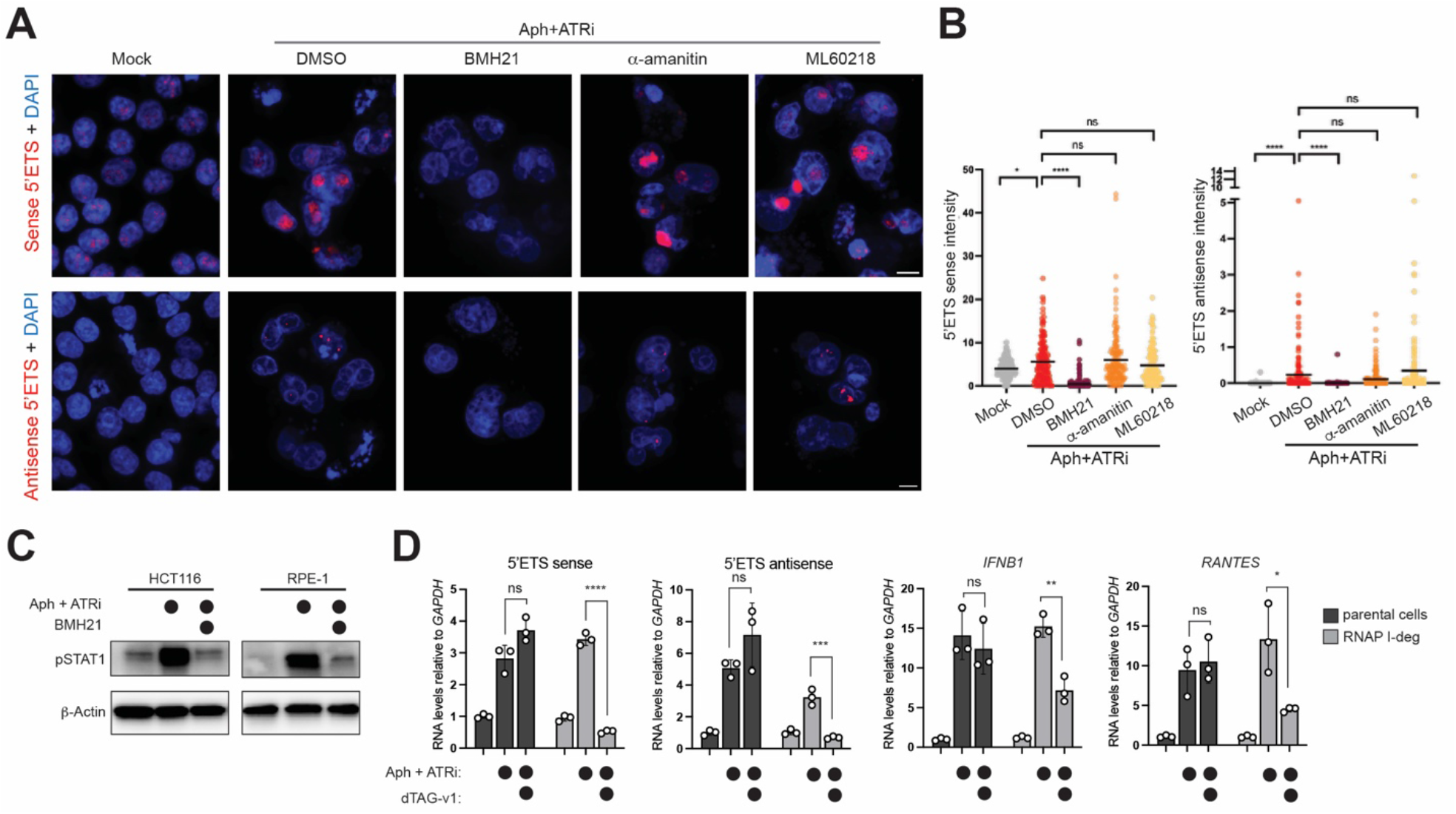
Replication stress-induced aberrant rDNA transcripts are generated by RNAP I and drive RLR activation. (A) RNA-FISH analysis of sense and antisense 5′ETS RNA in HCT116 cells under mock or Aph+ATRi treatment (48 h). Inhibitors of rDNA transcription (BMH21, 0.5 μM), RNAP II (α-amanitin, 5 μM), and RNAP III (ML60218, 25 μM) were added 2 h prior to imaging. (B) Quantitation of RNA-FISH signal in (A). Each data point represents an individual cell (n=247, 179, 236, 152, 141 from left to right) from ~20 fields of view per condition. \*\*\*\**p* <0.0001,\**p* <0.05, not significant (ns) *p* > 0.05 by two-tailed Mann–Whitney test. (C) Western blot analysis of pSTAT1. HCT116 and RPE-1 cells were treated with Aph+ATRi with or without BMH21 for 48 h. (D) Effect of RNAP I degradation on RLR signaling. RT-qPCR analysis showing indicated RNA levels relative to *GAPDH* mRNA in the parental or RNAP I-deg RPE-1 cells with or without dTAG-v1 (0.5 μM). Values represent mean ± SD from 3 independent experiments. *P*-values were based on two-tailed student’s t test. \*\*\*\**p* <0.0001, \*\*\**p* <0.001, \*\**p* <0.01, \**p* <0.05, not significant (ns) *p* > 0.05

**Fig. S4.**
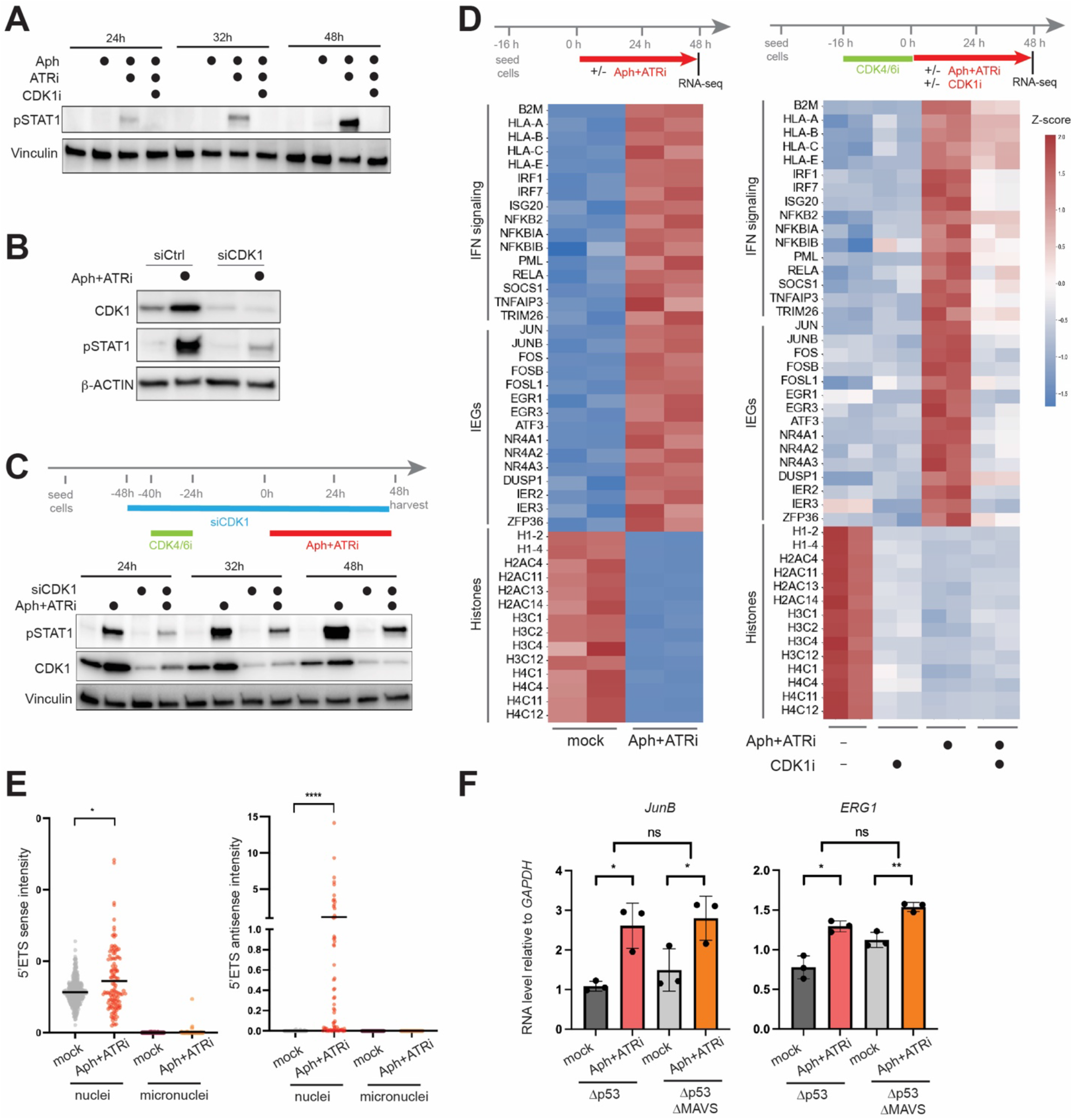
CDK1 amplifies the innate immune response to replication stress. (A) Western blot analysis of pSTAT1 in RPE-1 cells treated with Aph, Aph+ATRi, or Aph+ATRi+CDK1i for indicated duration. (B) Effect of CDK1 knockdown on innate immune response to Aph+ATRi. Western blot analysis of pSTAT1 and CDK1 in the presence of siRNA targeting CDK1 (siCDK1) or control siRNA (siCtrl). RPE-1 cells were treated with siRNA for 48 h and subjected to Aph+ATRi for additional 24 h. These blots are reproduced in Fig. 4C with additional blots. (C) Effect of siCDK1 on G1-synchronized cells. Cells were synchronized with CDK4/6i (palbociclib) prior to Aph+ATRi treatment as indicated in the schematic. Similar effect of siCDK1 was observed as in (B). (D) RNA-seq analysis of cellular response to replication stress in HCT116 cells. Left: Aph+ATRi treatment for 48 h induces IFN signaling genes and immediate early genes (IEGs), while suppressing histone genes. The histone suppression is consistent with G2 arrest. Right: Cells were synchronized in G1 with CDK4/6i and released into S phase in the presence of Aph+ATRi and/or CDK1i as indicated. Aph+ATRi again induced IFN signaling genes and IEGs while suppressing histone genes. CDK1i alone had little effect on IFN signaling or IEG induction, but suppressed histone gene expression, consistent with its known ability to induce G2 arrest. In the presence of Aph+ATRi, CDK1i suppressed IFN signaling genes and IEGs, with little additional effect on histone genes. These observations further suggest that the effect of CDK1 under replication stress is largely independent of its canonical role in cell-cycle progression. (E) RNA-FISH analysis of 5’ETS sense and antisense RNA from the main nuclei vs micronuclei. RNA-FISH intensity was measured from RPE-1 subjected to Aph+ATRi for 48 h. Each data point represents an individual cell (sense: n=469 and 122 for mock and Aph+ATRi; antisense: n=315 and 68 for mock and Aph+ATRi) from ~20 z-stacked fields of view per condition. *\*\*\*\*p<0*.*0001,*p<0*.*05*, by two-tailed Mann–Whitney test. (F) RT-qPCR analysis of IEGs in MAVS-sufficient vs. -deficient RPE-1 cells in the Δp53 background. *GAPDH* mRNA was used as an internal control.\**p* <0.05, \*\**p*<0.01, from Welch’s t-test.

**Fig. S5.**
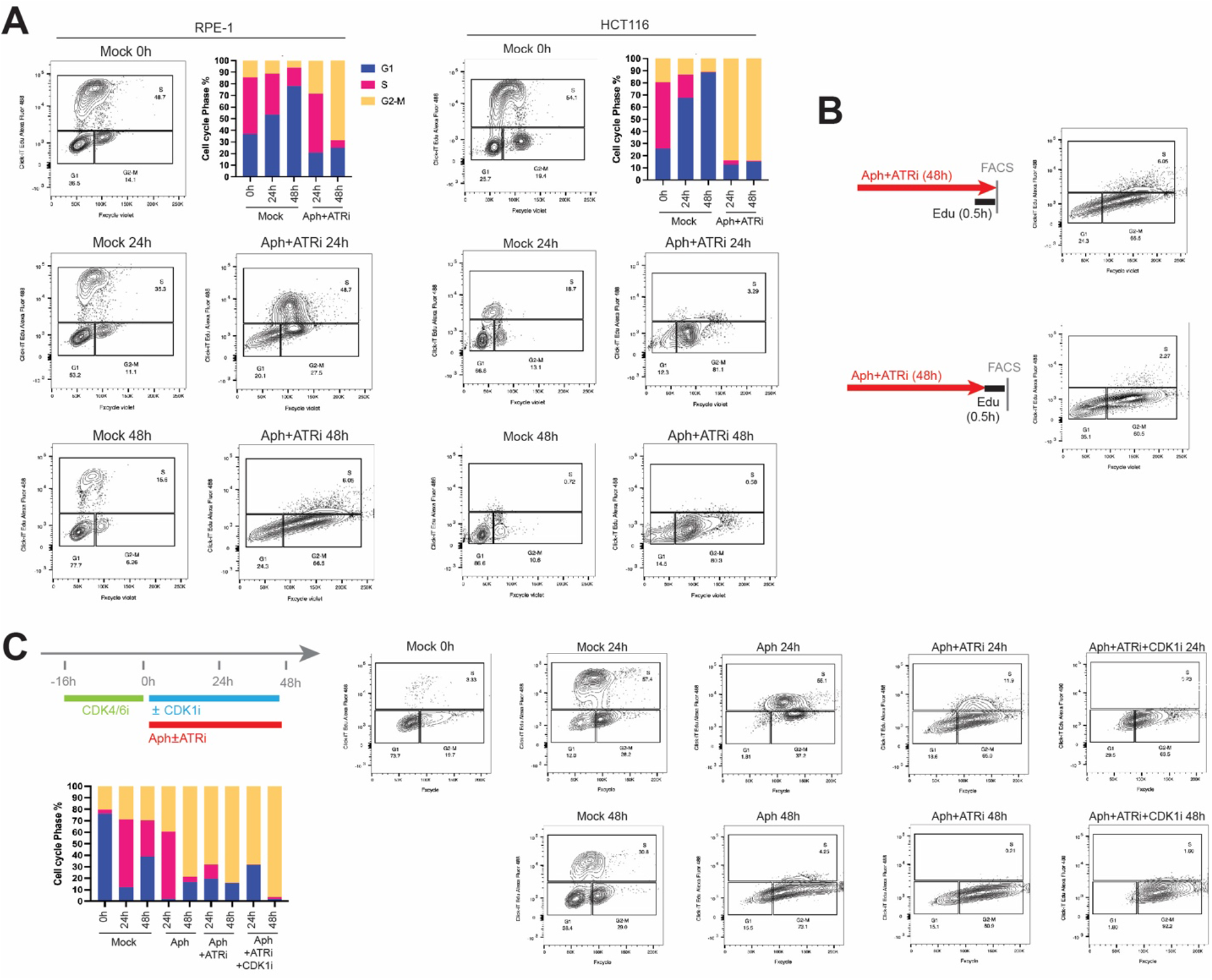
Replication stress leads to G2 arrest. (A) Cell cycle analysis by flow cytometry. RPE-1 and HCT116 cells were treated with Aph+ATRi for 0, 24, 48 h. FxCycle Violet stain and click it EdU–Alexa Fluor 488 were used to measure DNA content and DNA synthesis, respectively, and to define G1, S, and G2–M populations. Representative plots and quantification are shown. (B) EdU incorporation analysis by flow cytometry. Cells were treated with Aph+ATRi for 48 h and pulsed with EdU (0.5 h), during which Aph+ATRi was kept (top) or removed (bottom). Both methods show similar results. Unless mentioned otherwise, Aph+ATRi was kept during EdU labeling throughout this study. (C) Impact of ATRi and CDK1i on cell cycle phases during replication stress. Cells were treated with Aph, Aph+ATRi, or Aph+ATRi+CDK1i over the indicated timeline and analyzed as in (A).

**Fig. S6.**
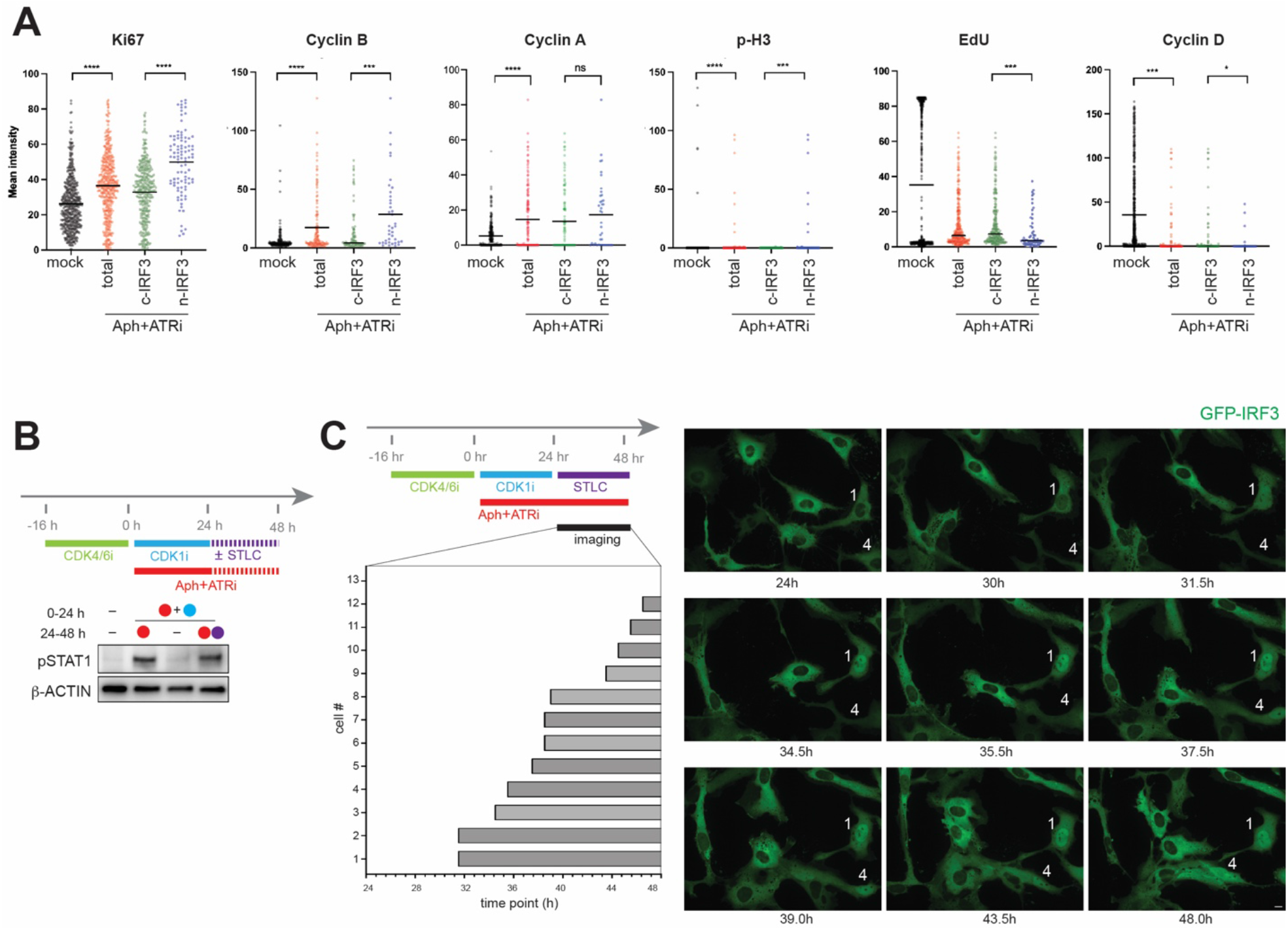
Replication stress induces RLR activation independent of mitosis. (A) Immunofluorescence quantification of cell cycle markers. Mean intensities of Ki67, Cyclin B, Cyclin A, p-H3, EdU, and Cyclin D are shown in mock and 32 h Aph+ATRi-treated RPE-1 cells, grouped by total, cytosolic IRF3 (c-IRF3), and nuclear IRF3 (n-IRF3) cells. Each data point indicates an individual cell. \*\*\*\**p* <0.0001, \*\*\**p* <0.001, \**p* <0.05, not significant (ns) *p* > 0.05, by two-tailed Mann–Whitney test. (B) Impact of STLC on RLR signaling as measured by pSTAT1 immunoblot. Cells were synchronized in G1 with CDK4/6i (palbociclib, 0.2 μM) and released into S phase in the presence of Aph+ATRi and CDK1i (RO3306, 6 μM). CDK1i was used to block G2>M transition. Upon CDK1i release, cells were maintained in Aph+ATRi for an additional 24 h, with or without the mitotic inhibitor STLC (2 μM). Analysis of pSTAT1 shows that continued Aph+ATRi in the presence of CDK1 activity is needed for RLR signaling, but this is independent of mitotic progression. (C) Time-lapsed imaging of GFP-IRF3 translocation. Top left: RPE-1 cells were treated as indicated. Bottom left: 12 cells showing IRF3 nuclear translocation were numbered in the order of IRF3 translocation. Gray bars indicate duration of nuclear GFP-IRF3. As expected from the known function of STLC in blocking mitosis, no cell underwent mitosis as monitored by cell rounding. Right: representative time-lapsed images of GFP-IRF3 (green) with cell identities indicated.

**Fig. S7.**
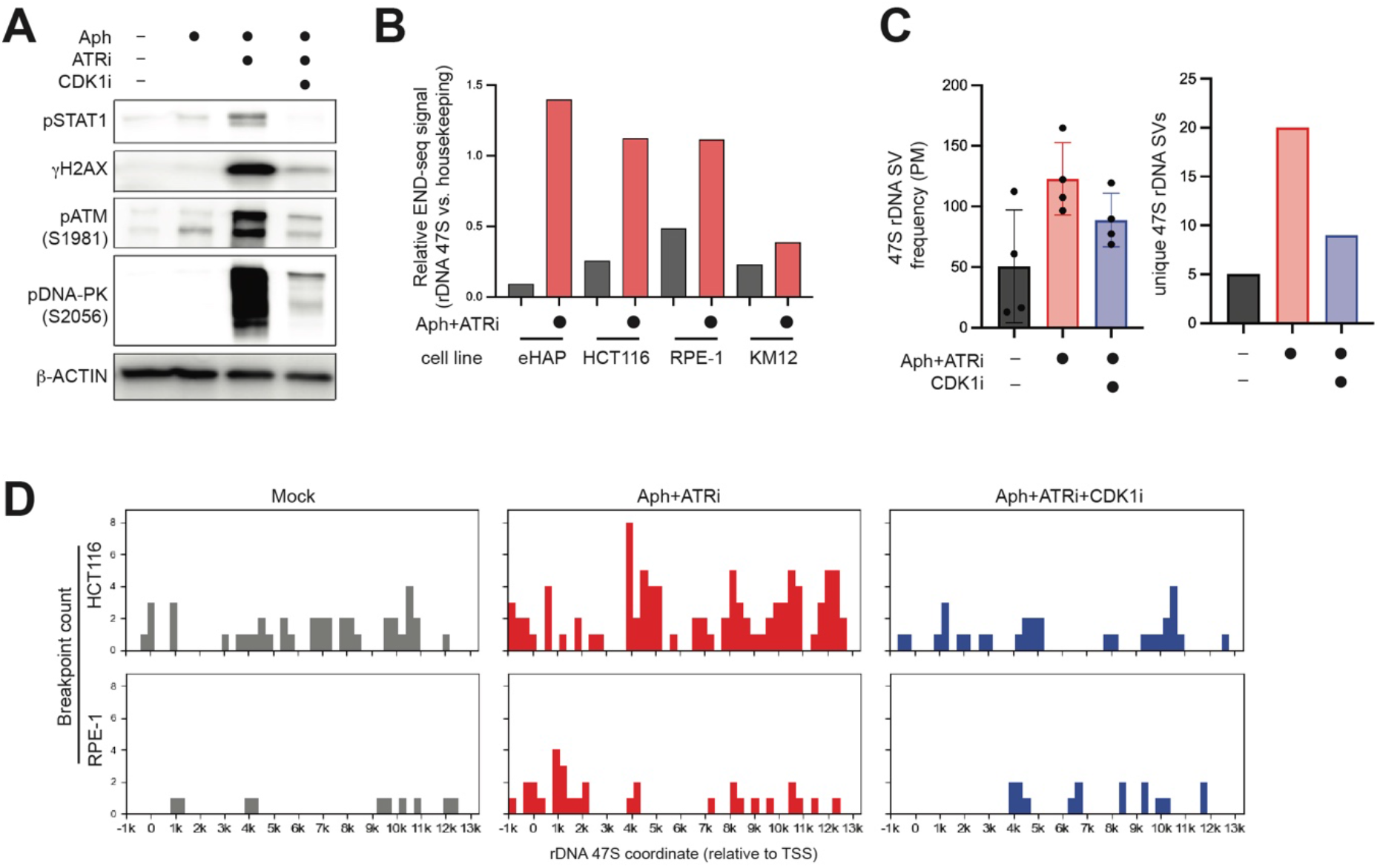
CDK1 activity promotes replication stress–induced rDNA instability. (A) Western blot analysis of DNA damage markers. HCT116 cells were treated with Aph, ATRi, and/or CDK1i for 32 prior to harvesting. pSTAT1, γH2AX, pATM (S1981), and pDNA-PK (S2056) are shown. (B) END-seq intensities in rDNA 47S region relative to housekeeping genes. END-seq signal was quantified as AUC (area under curve) for the 47S region and housekeeping genes (*GAPDH, ACTB, RPL3, RPS6, RPS12, RPS18*) in mock and Aph+ATRi-treated samples. The ratios of 47S to housekeeping gene signals are plotted. END-seq data were obtained from GSE149709. (C) Structural variations (SVs) in 47S rDNA induced by Aph+ATRi in the presence and absence of CDK1i. RPE-1 cells were treated with indicated drugs for 48 h and whole genome sequencing (WGS) was performed. Sequences were mapped to rDNA repeat unit (IGS-47S), followed by SV analysis using Delly. All SV types (deletions, inversions and duplications) were aggregated, and only SV events overlapping with the 47S region were retained for downstream analyses. Left: 47S SV frequency as measured by reads supporting 47S region SVs per million reads mapped to 47S region (PM). Right: unique SV events in 47S region. (D) Distribution of SV breakpoints across the 47S rDNA region and 1 kb upstream of the transcription start site (TSS; −1 kb) in HCT116 and RPE-1 cells treated with mock (grey), Aph+ATRi (red), or Aph+ATRi+CDK1i (blue). The number of SV breakpoints is plotted as a function of position relative to the 47S TSS in bins of 300 bp.

**Fig. S8.**
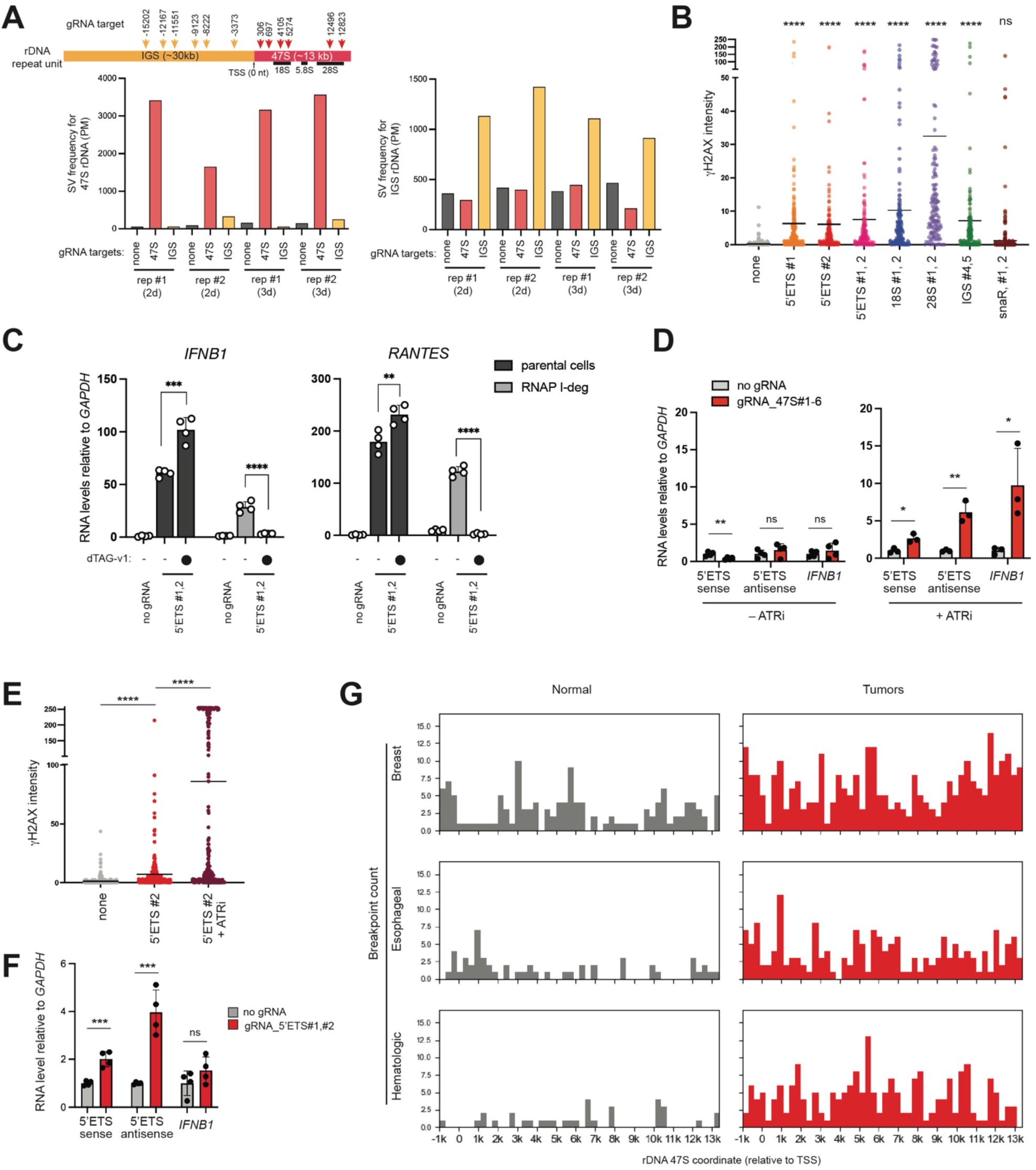
Replication stress compromises rDNA integrity and rDNA breaks are sufficient to activate antiviral signaling. (A) Cas9-induced structural variations in rDNA. HCT116 cells were electroporated with Cas9 complexed with 6 gRNAs targeting the IGS or 47S regions, as indicated in the schematic (top). Cells were harvested 48 h (2d) or 72 h (3d) post-electroporation for whole genome sequencing (WGS). Structural variation (SV) frequency in the 47S or IGS regions was calculated as reads supporting SVs normalized to total reads mapped to the respective region and expressed per million reads (PM). The results show that SVs reflect locus-specific DNA breaks; Cas9 targeting of the 47S selectively increased SVs within 47S, but not in the IGS regions, and vice versa. Individual biological repeats (rep #1, #2) are shown separately. (B) Immunofluorescence analysis of γH2AX. RPE-1 cells were analyzed 48 h post-electroporation with Cas9 and the indicated gRNAs. Each point represents an individual cell (n=296, 234, 176, 170, 192, 144, 184, 415 from left to right) from ~50 fields of view per condition. Statistical comparisons of each condition vs no gRNA control by two-tailed Mann-Whitney test. \*\*\*\**p*<0.0001; ns, not significant. (C) RT-qPCR analysis showing indicated RNA levels relative to *GAPDH* mRNA in the parental or RNAP I-deg RPE-1 cells with or without dTAG-v1 (0.5 μM) upon Cas9-induced breaks in 47S 5’ETS. Values represent mean ± SD from 4 technical replicates. *P*-values were based on two-tailed student’s t test. \*\*\*\**p* <0.0001, \*\*\**p* <0.001, \*\**p* <0.01. (D) Effect of ATRi on innate immune activation in response to Cas9-induced rDNA break. HCT116 cells were electroporated with Cas9 in complex with six gRNA targeting 47S (#1-6), treated with or without ATRi and harvested 48 h post-electroporation for RT-qPCR analysis. RNA levels were normalized to GAPDH and plotted relative to the corresponding no gRNA controls. \*\**p* <0.01, \**p* <0.05, not significant (ns) *p* > 0.05 from two-tailed student’s t test. (E) Immunofluorescence analysis of γH2AX. RPE-1 cells were analyzed 48 h post-electroporation with Cas9 and gRNA targeting 5’ETS #2 in the presence and absence of ATRi. Each point represents individual cell (n=366, 256, 244 from left to right) from ~50 fields of view per condition. \*\*\*\**p*<0.0001 by two-tailed Mann-Whitney test. (F) Effect of Cas9-induced rDNA break on induction of 5’ETS RNA and *IFNB* in MAVS KO HCT116 cells. Cells were electroporated with Cas9 in complex with 5’ETS (#1,2) in the presence of ATRi and harvested 48 h post-electroporation for RT-qPCR analysis. RNA levels were normalized to GAPDH and plotted relative to the corresponding no gRNA controls. \*\*\**p* <0.001, not significant (ns) *p* > 0.05 from two-tailed student’s t test. (G) Distribution of SV breakpoints across the 47S rDNA region and 1 kb upstream of the transcription start site (TSS; −1 kb) in breast, esophageal and hematologic tumors (red) and matched normal samples (grey) from Fig. 4I. Data were analyzed as in (D).

**Table S1.**
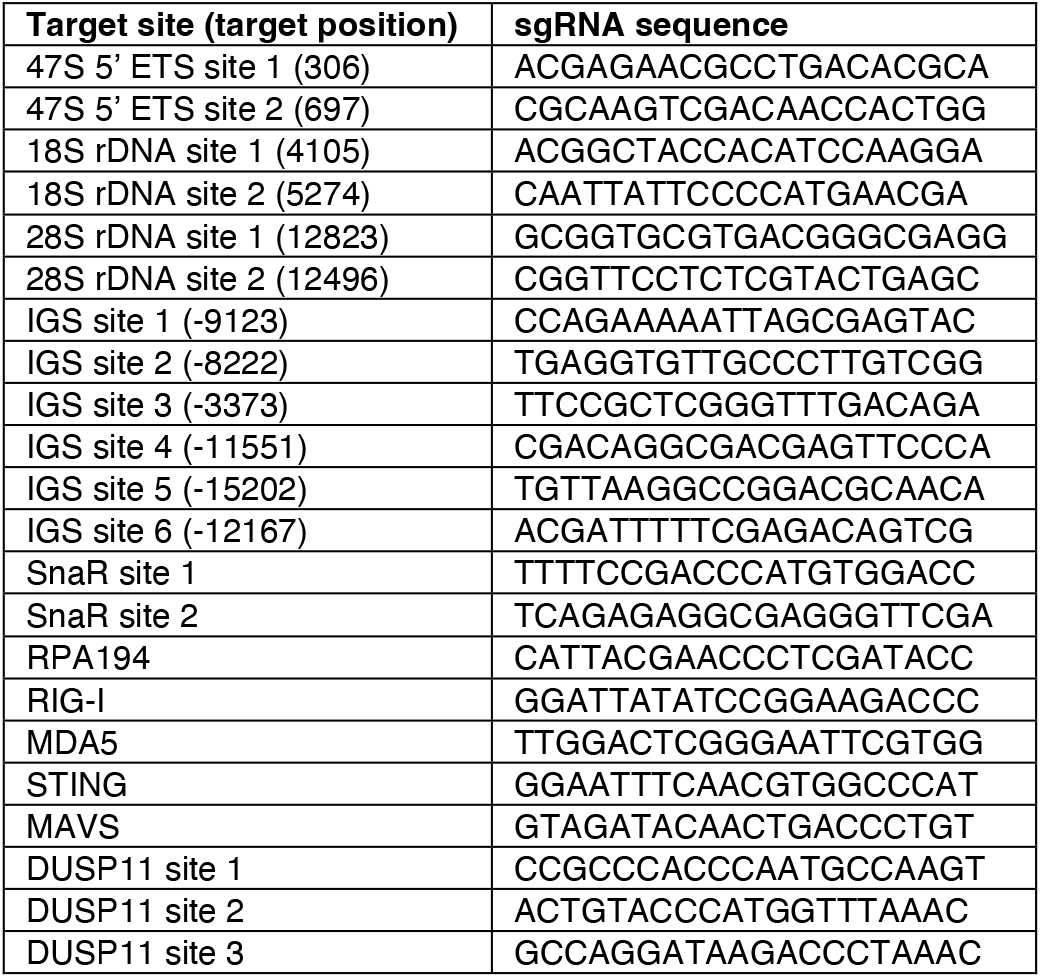
sgRNA sequences for CRISPR-Cas9.

**Table S2.**
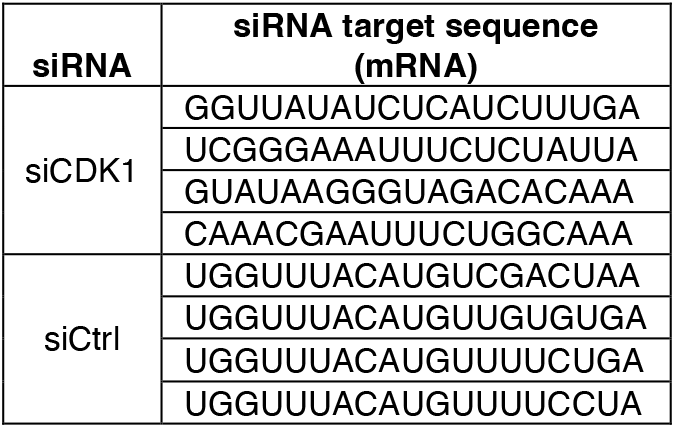
siRNA target sequences.

**Table S3.**
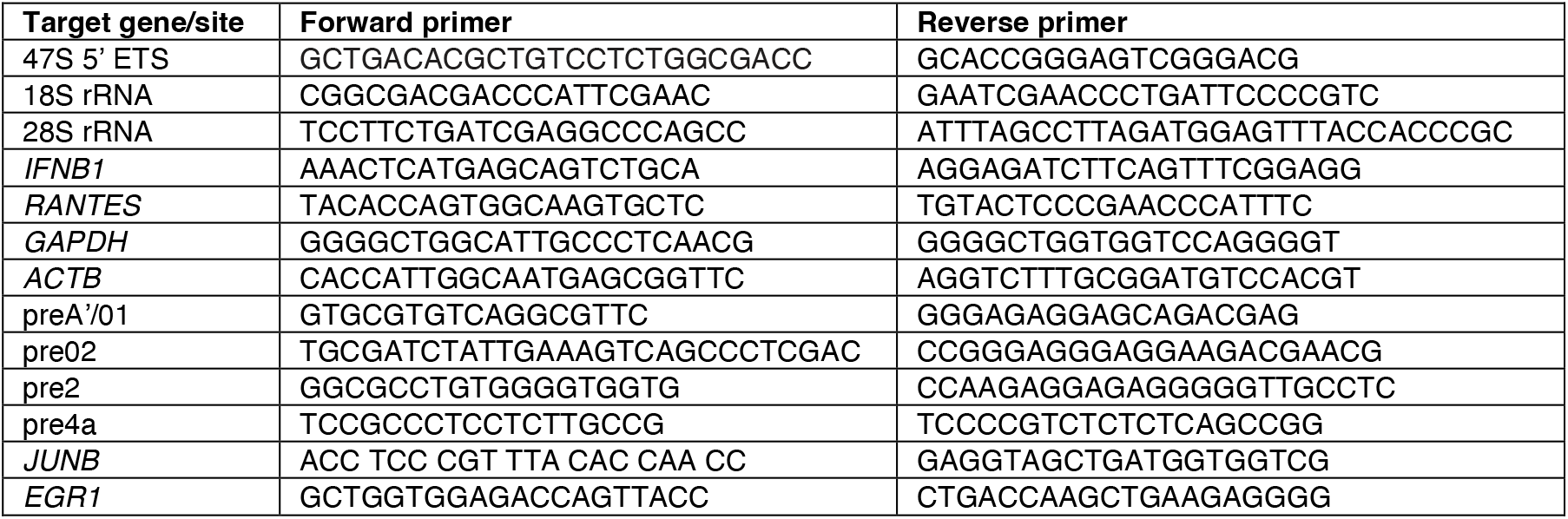
qPCR primer sequences.

**Table S4.**
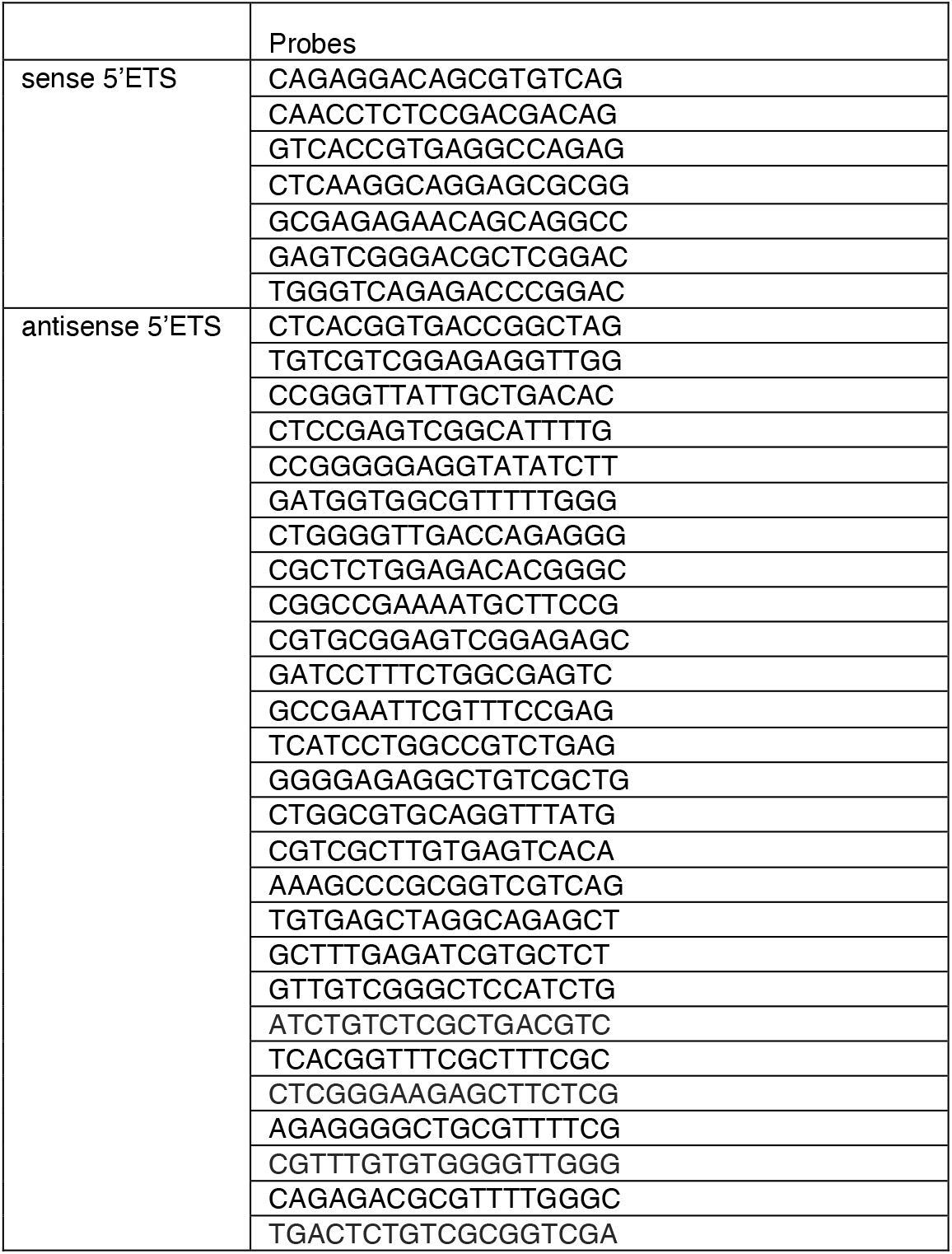
Stellaris RNA-FISH probes for detection of the sense and antisense 47S 5’ETS.

**Table S5.** Chemicals.

| Chemicals | Brand | Cat. Number |
| --- | --- | --- |
| Aphidicolin | Sigma | A0781 |
| VE-821 | selleckchem | S8007 |
| BMH-21 | selleckchem | S7718 |
| a-Amanitin | Tocris (R&D) | 4025 |
| ML-60218 | medchemexpress | HY-122122 |
| Palbociclib | selleckchem | S1116 |
| RO3306 | selleckchem | S7747 |
| dTAG-v1 | Tocris | 6914/5 |
| S-Trityl-L-cysteine (STLC) | medchemexpress | HY-W011102 |

